# A data-driven algorithm to determine ^1^H-MRS basis set composition

**DOI:** 10.1101/2024.09.11.612503

**Authors:** Christopher W. Davies-Jenkins, Helge J. Zöllner, Dunja Simicic, Seyma Alcicek, Richard A.E. Edden, Georg Oeltzschner

## Abstract

**Purpose:** Metabolite amplitude estimates derived from linear combination modeling of MR spectra depend upon the precise list of constituent metabolite basis functions used (the “basis set”). The absence of clear consensus on the “ideal” composition or objective criteria to determine the suitability of a particular basis set contributes to the poor reproducibility of MRS. In this proof-of-concept study, we demonstrate a novel, data-driven approach for deciding the basis-set composition using Akaike information criteria (AIC).

**Methods:** We have developed an algorithm that iteratively adds metabolites to the basis set using iterative modeling, informed by AIC scores. We investigated two quantitative “stopping conditions”, referred to as max-AIC and zero-amplitude, and whether to optimize the selection of basis set on a per-spectrum basis or at the group level. The algorithm was tested using two groups of synthetic in-vivo-like spectra representing healthy brain and tumor spectra, respectively, and the derived basis sets (and metabolite amplitude estimates) were compared to the ground truth.

**Results:** All derived basis sets correctly identified high-concentration metabolites and provided reasonable fits of the spectra. At the single-spectrum level, the two stopping conditions derived the underlying basis set with 84-88% accuracy. When optimizing across a group, basis set determination accuracy improved to 89-92%.

**Conclusion:** Data-driven determination of the basis set composition is feasible. With refinement, this approach could provide a valuable data-driven way to derive or refine basis sets, reducing the operator bias of MRS analyses, enhancing the objectivity of quantitative analyses, and increasing the clinical viability of MRS.

## 1. Introduction

In-vivo proton magnetic resonance spectroscopy (MRS) is a non-invasive method for estimating the concentrations of approximately 20 metabolites in the human brain. Levels of neurotransmitters and antioxidants, for example, may serve as biomarkers of function and pathology, but also general indicators of neuronal health and cell proliferation. Extracting metabolite concentrations from the in-vivo spectrum depends on reliably resolving the contributions of constituent signals. For this task, expert consensus recommends^1,2^ linear-combination modeling (LCM), a well-established method that uses a weighted sum of simulated signals—*basis functions*—to approximate the measured spectrum.

Poor spectral dispersion at clinical field strengths causes overlap between metabolite signals, preventing them from being estimated independently. To achieve optimal precision and accuracy, the basis set included in LCM should be selected without bias. Including too few metabolites results in unmodeled or poorly modeled peaks. Most LCM implementations will attempt to reconcile the model by modulating the baseline and lineshape, or shifting metabolites to reduce the residual. Conversely, including too many basis functions in the model may result in overfitting the data—with noise or lineshape effects being incorrectly attributed to metabolite signals. Many compounds are present below the threshold of detection in the healthy brain, only reaching detectable levels in pathology (e.g., 2-hydroxyglutarate in primary brain tumors with isocitrate dehydrogenase mutations^3^).

In practice, choosing which metabolites to include in the basis set is delicate because *many different* compositions are *admissible*, i.e., are reasonable choices. Several fundamental challenges arise:

1. Wrongly in- or ex-cluding basis functions can lead to substantial biases of^4–8^ and interactions between^9,10^ metabolite amplitude estimates.
2. There is no consensus on the ideal basis set composition even for the healthy brain, resulting in significant analytic variability between research groups.
3. Defining basis-set composition currently requires a-priori knowledge (e.g., an external brain tumor diagnosis to adequately model a brain tumor spectrum), limiting the clinical application of MRS.
4. The number of metabolites that can be modeled with reasonable accuracy and precision depends strongly on the quality of the data. High-SNR, well-resolved spectra allow the modeling of metabolites that might otherwise have simply led to the overfitting of lower-quality data.
5. Finally, no objective criteria exist to assess whether a specific basis set will overfit, underfit, or appropriately model a spectrum.

To address these challenges, we investigated the feasibility of determining the appropriate basis set composition directly from the spectrum itself using model selection. Model selection is the process of determining the most suitable model from a list of potential candidates informed by a quantitative information criterion (IC). There are several specific definitions of ICs^11–14^, but generally, they attempt to “score” candidate models, balancing goodness-of-fit and model complexity in a single value^15,16^. Higher IC scores are interpreted as a proxy measure of model parsimony, i.e., the “cost-effectiveness” of particular candidate models; ICs have previously been used in MRS to derive optimal baseline model parameters^17^.

In this proof-of-concept study, we demonstrate a novel automated data-driven procedure—using iterative fitting of the spectra informed by IC scores—to determine the appropriate composition of the basis set. We developed two variations of this algorithm: the first determines the optimal basis set for a single spectrum, and the second for a group of spectra. We also designed two different stopping conditions: the first stops adding metabolites to the basis set after reaching the maximum IC score, and the second keeps adding metabolites beyond this point until they no longer contribute any signal to the fit. We tested the performance of the algorithm and its variations in three distinct scenarios. Firstly, we generated two distinct classes of in-vivo-like simulated spectra, representing high-quality spectra from the healthy brain and low-grade glioma, respectively. The latter included two oncometabolites that were not present in the healthy brain data. We quantified how overlap between the algorithm-selected basis sets and the ground-truth basis sets from which spectra were generated. Secondly, we probed the limitations of our algorithm by adding varying degrees of noise and linebroadening to the synthetic healthy brain spectra and, as before, examined where the algorithm-selected basis sets began to deviate from the ground truth. Finally, we applied the algorithm to real brain spectra.

## 2. Methods

### 2.1. Synthetic in-vivo-like data generation

We first simulated realistic in-vivo-like spectra to establish a known ground truth. We generated two datasets of 100 spectra, each with distinct spectral characteristics: healthy-appearing brain spectra and those typically seen in low-grade glioma, respectively. We chose the latter case as these spectra commonly feature metabolites that are effectively absent from healthy brain tissue (cystathionine^18,19^ and 2-hydroxyglutarate^20^) and exhibit markedly different amplitudes of major metabolites.

Synthetic spectra were derived from a “library set” of metabolite basis functions, simulated using the density-matrix formalism of a 2D-localized 101 x 101 spatial grid (field of view 50% larger than the voxel) implemented in MRSCloud^21^, which is based on FID-A^22^. We synthesized 3T sLASER spectra to reflect a typical acquisition used to measure 2HG (TE = 97 ms with TE1/2 = 32/65 ms; 8192 complex points (downsampled to 2048 for the synthetic data); spectral width 4000 Hz).

We simulated 32 metabolite basis functions: Acetoacetate, AcAc; acetate, Ace; alanine, Ala; ascorbate, Asc; aspartate, Asp; citrate, Cit; creatine, Cr; creatine methylene, CrCH2; cystathionine, Cystat; ethanolamine, EA; ethanol, EtOH; γ-aminobutyric acid, GABA; Glycerophosphocholine, GPC; glutathione, GSH; glucose, Glc; glutamine, Gln; glutamate, Glu; glycine, Gly; myo-inositol, mI; lactate, Lac; N-acetylaspartate, NAA; N-acetylaspartylglutamate, NAAG; phosphocholine, PCh; phosphocreatine, PCr; phosphoethanolamine, PE; phenylalanine, Phenyl; scyllo-inositol, sI; serine, Ser; taurine, Tau; tyrosine, Tyros; β-hydroxybutyrate, bHB; and 2-hydroxyglutarate, 2HG. We also included basis functions for 5 macromolecular (MM09, MM12, MM14, MM17, MM20) and 3 lipid resonances (Lip09, Lip13, Lip20). Out of the full set of 40 basis functions, we selected 26 to assemble the synthetic healthy ground-truth spectra: Asc; Asp; Cr; CrCH2; GABA; GPC; GSH; Gln; Glu; mI; Lac; NAA; NAAG; PCh; PCr; PE; sI; Tau; MM09; MM12; MM14; MM17; MM20; Lip09; Lip13; and Lip20. The locations, widths, and relative amplitudes of the parameterized lipid and MM resonances are reported in **Table S1** and the full library is visualized in **Figure S1**.

For the tumor spectra, we added additional contributions from 2HG and Cystat for a total of 28 basis functions in the tumor ground-truth basis set. The Osprey function “OspreyGenerateSpectra”^23^ was used to combine the individual simulated profiles with metabolite-specific amplitudes, Gaussian and Lorentzian linebroadening terms, and white noise, taken from Gaussian distributions. Crucially, each group was defined with distinct model parameter ranges, which were informed by previously modeled in-vivo data^7,24^. Briefly, besides the additional 2HG and Cystat contributions, the tumor spectra also had higher tCho (100%), lower tNAA (60%), higher mI (30%), lower Glu (25%), higher Lac (1000%), higher lip09 (550%), and higher Lip20 (550%). The means and standard deviations of the parameters for each group are fully reported in **Table S2** and the spectra are visualized in **Figure S2.** The reported FWHM and SNR values were calculated using the FID-A functions “op_getLW” and “op_getSNR”, respectively. The FWHM was calculated on the 3-ppm Cr peak by taking the average of a direct FWHM measurement and Lorentzian model. The SNR is calculated in the frequency domain by taking the ratio of the maximum of the NAA peak to the standard deviation of a signal-free region (between –2 and –1 ppm) following the removal of a linear baseline.

### 2.2. Basis set determination algorithm

Our proposed algorithm uses IC scores to build the basis set in an iterative process. Introducing more basis functions reduces model residuals but will add model parameters (i.e. more metabolite-specific amplitudes, frequency shifts, and lineshape parameters). Determining the IC scores for each potential basis set composition allows us to counterbalance these two competing aspects to arrive at a parsimonious compromise, i.e., maximize the goodness-of-fit without overfitting.

#### 2.2.1. Information criteria

We implemented three IC definitions within our LCM framework: the Akaike IC (AIC), corrected AIC (AICc), and Bayesian IC (BIC). They differ in how they regularize model complexity against goodness-of-fit. There was unanimous agreement between the IC scores on the order in which metabolite basis functions were prioritized (**Figure S3**). However, initial testing revealed that the basis sets derived by the BIC scores were too sparse compared to those of the AIC and AICc. We elected to proceed using the AIC as the primary model-performance metric in the algorithm. The IC are defined as follows:

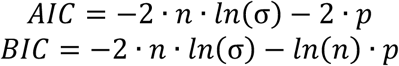

where *n* is the number of points, *p* is the total number of model parameters, and 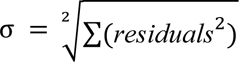, the root sum square of the fit residual.

#### 2.2.2. Algorithm implementation

We designed an algorithm to determine which of the library set of 40 basis functions (32 metabolites plus 8 MM/Lips) merits inclusion in the “selected set”. The selected set (which is initially empty) is built up one basis function at a time—the function added in each round of the algorithm is chosen from among the remaining candidates in the library set, based upon which model had the highest AIC. Thus, the algorithm consists of two nested loops: the outer loop (of rounds) to determine which is the next candidate to add to the selected set, and the inner loop (of steps) which consists of modeling the data with the current selected set plus one additional candidate.

Both loops fill the selected basis set iteratively until a certain stopping condition is triggered. The algorithm is illustrated in **Figure 1A**, and a **Supplementary Video**, with the specific steps as follows:

1. Initialize an empty selected set and a full library set.
2. Perform a preliminary fit using the full library set to estimate global lineshape and phase parameters which are then fixed during subsequent model calls.
3. Round 1:

a. Step 1 consists of modeling the spectrum with a candidate set containing only the first library basis function.
b. The remaining 39 steps of Round 1 consist of modeling the spectrum with a candidate set consisting of each library basis function. The AIC is calculated for each model.
c. Round 1 concludes by moving the candidate function that resulted in the model with the highest AIC into the selected basis set.
4. The *N*-th Round then steps over the (41 − *N*) remaining library functions, thus:

a. Form the candidate basis set from the selected set and the *n*^th^ basis function from the remaining library set.
b. Model the data with the candidate set.
c. Calculate the AIC and note the amplitude of the *n*^th^ basis function within the model.
5. Check whether the stopping condition has been met (as described below).
6. If the stopping condition is not met, move the candidate basis function with the highest AIC score from the library set to the selected set and return to step 4.

**Figure 1.**
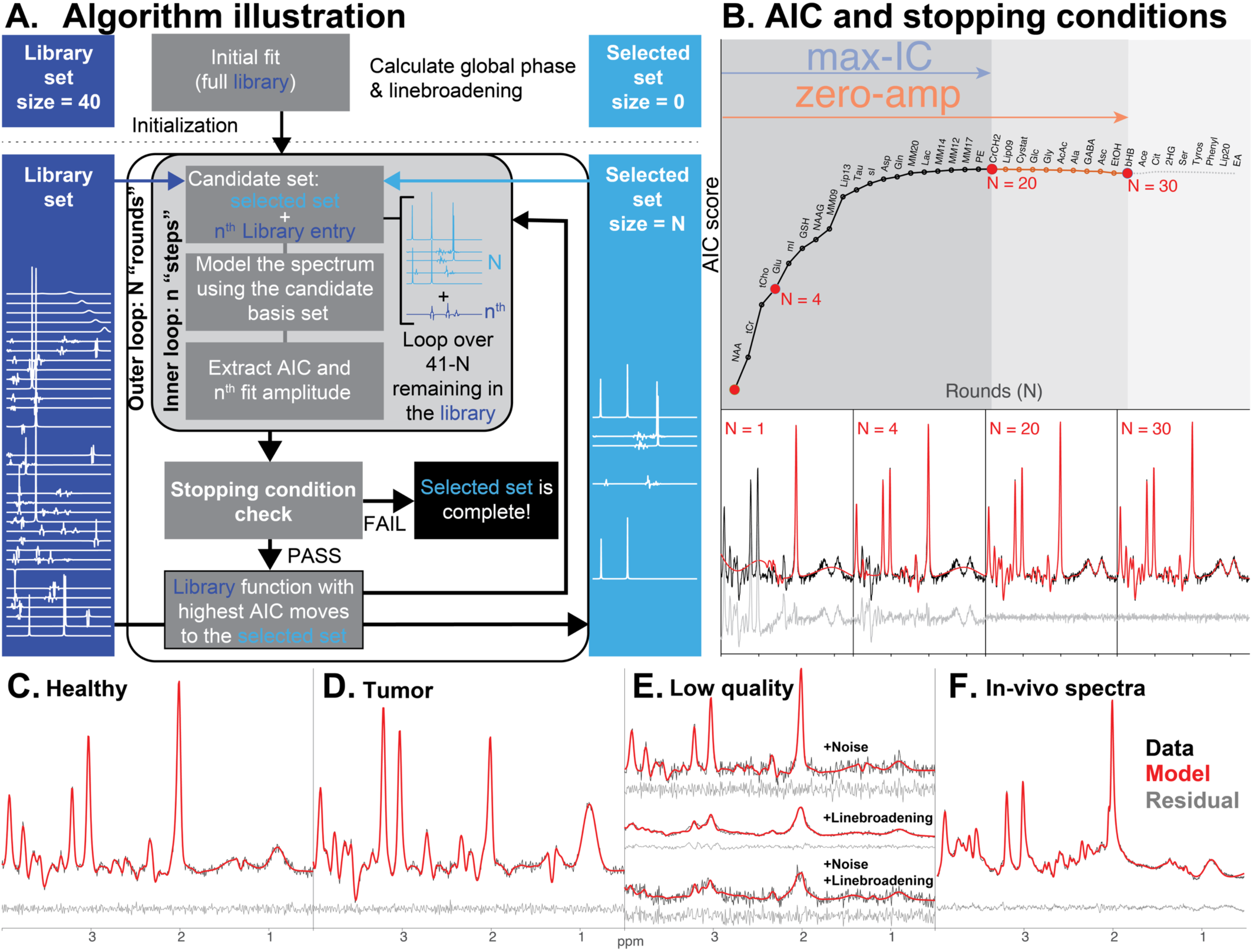
Basis set selection algorithm. **A.** A visualization of the algorithm. Metabolites are iteratively moved from a library set (dark blue) to a selected set (light blue) using AIC scores. The basis function that yields the best AIC score is moved to the selected set (unless a stopping condition is triggered). **B.** A visualization of the evolution of AIC scores as the basis set is optimized. The two stopping conditions are represented by the colored arrows and shaded regions. 4 example fits are shown in the panels below (corresponding to red points on the curve). **C–F.** Panels show representative spectra (black) and a corresponding model (red) derived from single-spectrum optimization and the zero-amplitude stopping condition. Model residuals are shown below (grey). **C.** A synthetic healthy-appearing spectrum **D.** A synthetic low-grade glioma spectrum **E.** Examples of synthetic lower-quality spectra. **F.** A representative real in-vivo spectrum.

We investigated two domains of selection, i.e. whether to select a basis set for each spectrum individually or to select one basis set at the group level for a set of spectra. For each domain, we also investigated two stopping conditions, i.e., when to stop adding basis functions to the selected set, see **Figure 1B**. The optimization domains are referred to as:

1. **“Single-spectrum”**: Executes the algorithm on each spectrum in isolation. Candidate basis functions are added based on the AIC scores derived from that spectrum alone, resulting in a spectrum-specific selected set.
2. **“Group-level”:** Executes the algorithm as described, but aggregates AIC scores across a *group* of spectra. At each step, we fit *all* spectra within a group with the candidate set and then use the median AIC score across the group to decide which metabolite is selected. This procedure results in a single basis set for the entire group.

The two stopping conditions were:

1. **“Max-AIC”:** The algorithm stops adding metabolites to the selected set once the AIC is decreased by the inclusion of the next basis function. For the group-level optimization, it stops once the *median* AIC across all datasets decreases.
2. **“Zero amplitude”:** The algorithm keeps adding metabolites until the round when all remaining candidate functions are modeled with zero amplitude. For the group-level optimization, it stops once all potential candidate functions are estimated with a *median* amplitude of zero across all datasets.

To stabilize the procedure, we instructed the algorithm to “link” certain pairs of practically indistinguishable basis functions, i.e., add both of them to the temporary candidate basis set as a single step, and, if selected, to add both to the selected basis set. Specifically, we linked Cr and PCr (referred to as the tCr step) and PCho and GPC (the tCho step).

#### 2.2.3. MRS modeling

All MRS modeling steps were performed using Osprey’s^24^ generalized LCM framework, introduced in our recent work^23,25^. Aside from the Gaussian and zero-order phase parameters—which were fixed following the initial modeling step—the model was defined using default in-vivo parameters, naïve to the composition of the simulated spectra. We performed optimization over the range of 0.5–4.2 ppm and included a non-regularized spline baseline with knot spacing of 0.5 ppm. Metabolite-specific Lorentzian linewidth and frequency shift parameters were regularized (as described in the original LCModel publication^26^) with expectation and standard deviation values of 2.75 ± 1.5 and 0 ± 3 Hz, respectively.

### 2.3. Statistical analysis

The algorithm’s performance was primarily judged by its ability to correctly identify the metabolite signals that were present in the synthetic spectra. This was assessed using three related metrics to compare each derived basis set to the ground truth set:

1. False positives: The number of basis functions selected for inclusion by the algorithm that were not present in the synthetic spectra (incorrectly included).
2. False negatives: The number of basis functions not selected for inclusion by the algorithm that were present in the synthetic spectra (incorrectly omitted).
3. The Sorensen-Dice coefficient (SDC): The SDC was used as a measure of overlap with the ground truth basis set, defined: 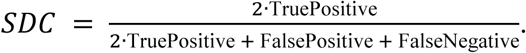

Finally, we investigated the effect that the derived basis set composition had on metabolite amplitude estimates. Amplitude deviations from the ground truth are reported as percentage changes relative to the known ground-truth value that entered the simulation. We also compared the amplitudes derived from each basis set to those derived from a fit using the *correct* ground truth basis set, i.e., modeling the data with only the metabolites we know to be present in the simulation. Distributions were compared using paired t-tests. All statistical analyses were conducted in Matlab 2024a.

### 2.4. Validation against fixed basis set compositions

To further demonstrate and validate our algorithm, we compared the amplitude estimates of the best-performing optimization approach (group-wise; zero-amplitude) to those derived from models using 4 expert-derived basis set compositions. These static basis set definitions were used to represent the nature and breadth of current practices in the field. We surveyed MRS experts and asked them to provide the composition of the basis set they would typically use for the analysis of healthy brain spectra. We selected four random respondents to the survey and we modeled the spectra using their suggested basis set composition. The 4 basis set compositions are detailed in **Table S3**.

### 2.5. Robustness to low SNR and high linewidth

To examine how our algorithm performed when spectral quality was reduced, we conducted a further analysis of the synthetic spectra. We augmented the healthy-brain dataset by introducing varying degrees of additional noise and Lorentzian linebroadening for a total of 9 data quality scenarios, each containing 100 spectra (**Figure S5**). These datasets were analyzed as described above, and the impact of linewidth and SNR was explored. Full implementation details are provided in the supplementary material.

### 2.6. Application to in-vivo data

To test how our algorithm performed in real spectra, we applied it to a freely available repository of 101 short-TE brain spectra^27^. In-vivo data were processed in Osprey^24^, and a new appropriately-simulated library set was generated^21^ which contained the same 40 metabolites as the synthetic data analysis. We ran all procedural variations of our algorithm to derive basis sets for these spectra. Full implementation details are provided in the supplementary material.

The algorithm implementation, specific Osprey version, and other code used to generate the results of this manuscript are available online (DOI: 10.17605/OSF.IO/P2USJ).

## 3. Results

### 3.1. Algorithmic trends of IC scores

As the algorithm added basis functions to the selected set, AIC scores followed a characteristic trajectory. High-concentration singlet resonances were initially prioritized by the algorithm, rapidly reducing model residuals and, consequently, increasing AIC scores. After this—as residual contributions reduced—the ascent of the AIC scores tapered off to a maximum (triggering the max-AIC stopping condition). According to the interpretation of AIC scores, this model is optimally parsimonious, and while additional basis functions *do* continue to reduce the residuals, they do not reduce them sufficiently to warrant the additional model parameters. This ascent of the AIC scores reliably identified the high-concentration metabolites that were present in the simulation.

Beyond the peak, AIC scores decreased as relatively smaller spectral contributions lowered the model’s cost-effectiveness. Eventually, the candidate metabolites contribute an amplitude of zero when added to the model. In other words, it costs model parameters but provides no benefit to the residual (triggering the 2^nd^ stopping condition: zero-amplitude). After this point, the remaining basis functions bring a continued decline. Importantly, ground-truth metabolites (those present in the synthetic spectra) tended to be included before the algorithm reached this final zone. Typical AIC score trajectories are illustrated in **Figure 2**.

**Figure 2.**
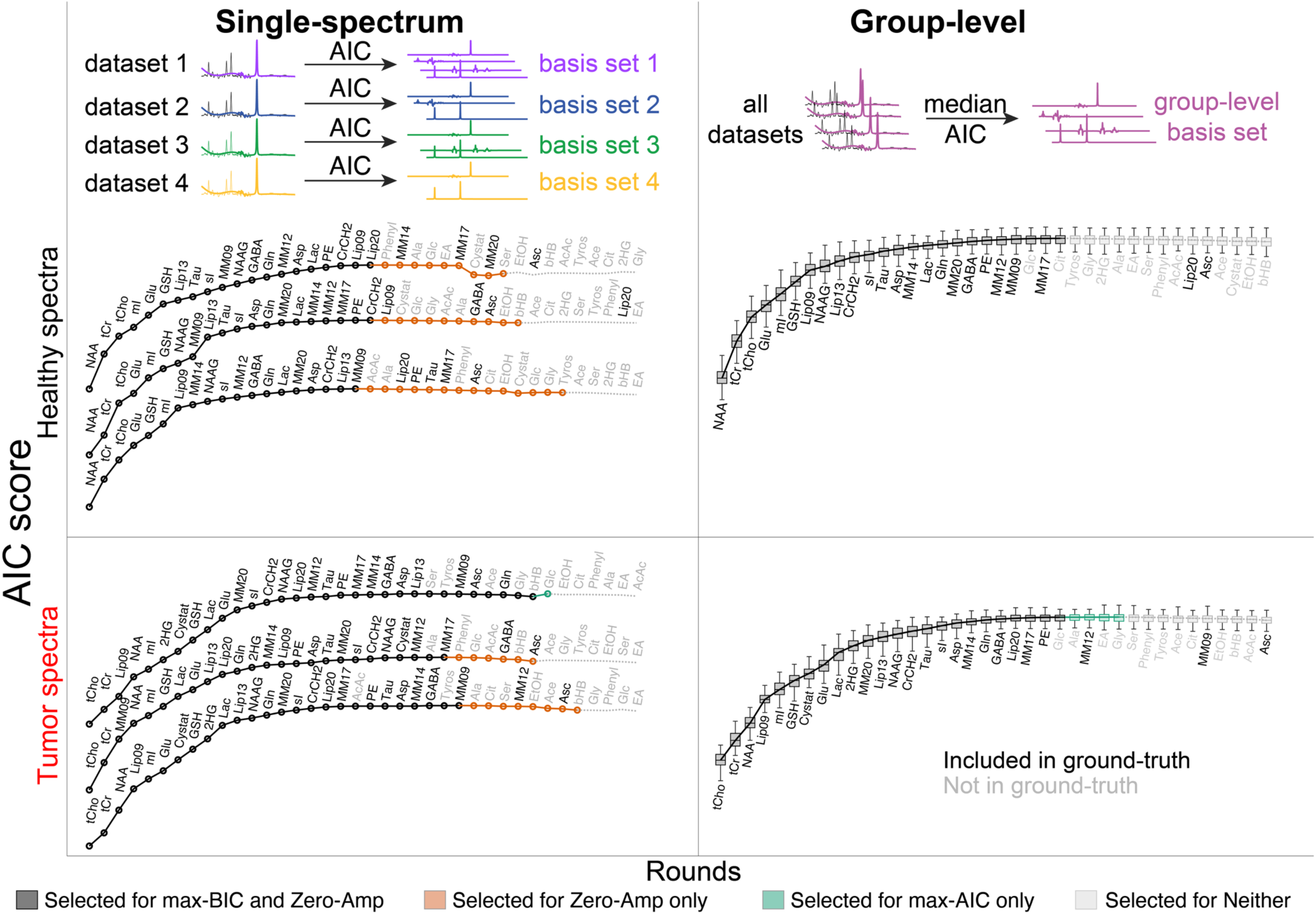
AIC scores as a function of algorithm round. The top row shows a conceptual visualization of the difference between the two optimization methods (single-spectrum vs. group-level). The lower two rows show the results for the healthy spectra (middle row) and tumor spectra (bottom row). The left column shows 3 (out of 100) example AIC curves for the single-subject optimizations, and the right column shows the group-level optimization, with the AIC distributions across the group shown. Text labels indicate the basis function added at each round, with label color reflecting the presence/absence (black/grey) of that metabolite in the ground-truth simulation. Line style is used to illustrate the extent of the derived basis sets in each stopping condition: included in both stopping conditions (solid black line), included only in the zero-amplitude condition (orange line), included in only the max-AIC condition (green), or included in neither (light-grey).

Figure 2 illustrates the between-spectrum variation of the basis set composition, and we have attempted to summarize the algorithmic trends of the single-spectrum optimization in Figure 3 and **Figure S4**. For healthy spectra, there is a clear prioritization of NAA, tCr, and tCho, in that order. The tumor spectra have a less clearly defined hierarchy of signals. The lower NAA and higher tCho amplitudes lead to a less unanimous prioritization of the singlets, and the larger lipid signals are chosen as early as the 2^nd^ round. Beyond the 5^th^ round, both groups of spectra had a diverse set of metabolites selected.

**Figure 3.**
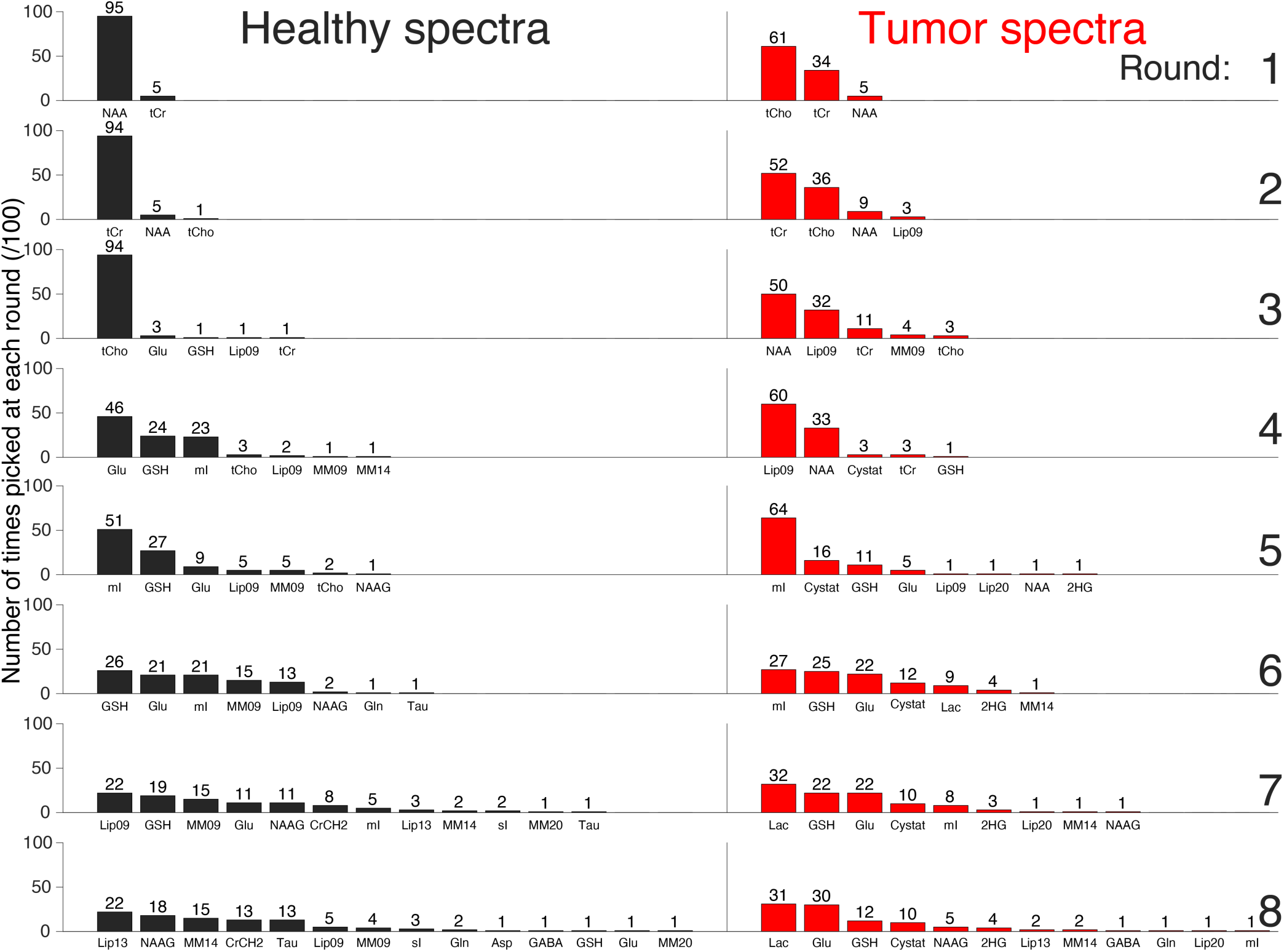
Histograms of the frequency with which particular metabolites were picked at each round of the algorithm. Rows represent the different rounds (increasing from top-to-bottom), and the two columns represent the two groups of spectra.

### 3.2. Overlap with the ground-truth basis set

For the single-spectrum optimization, the max-AIC stopping condition tended to be too conservative (few false positives, many false negatives) whereas the zero-amplitude condition tended to be too aggressive (few false negatives, many false positives). Bar plots showing the false positives, false negatives, and SDC are shown in Figure 4. Overall, the max-AIC stopping condition exhibited a better overlap with the ground-truth set, as evidenced by the SDC for both healthy-appearing spectra (mean difference = 5.0%, *p* = 0.014) and tumor spectra (mean difference = 4.7%, *p* = 0.010).

**Figure 4.**
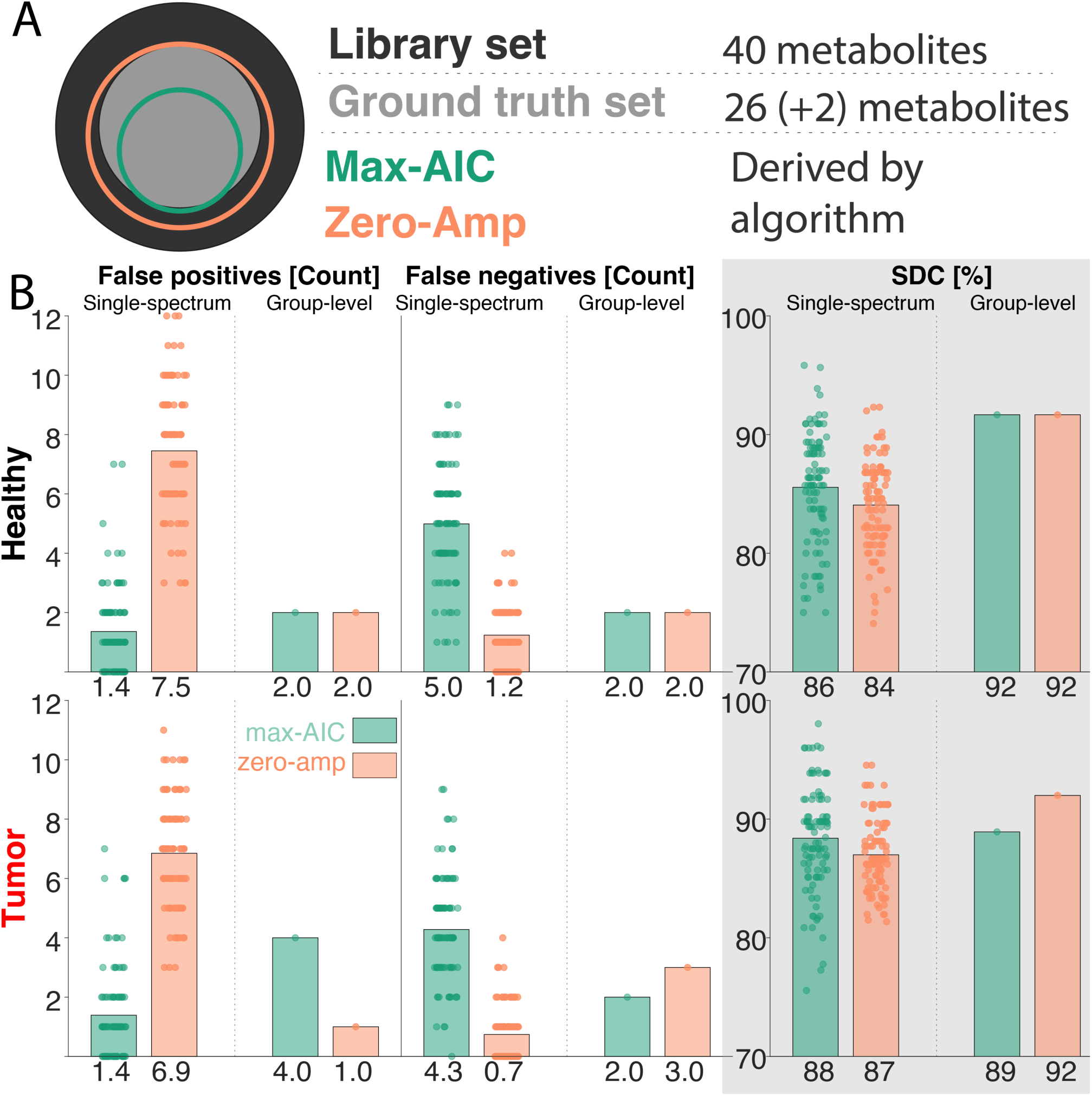
**A.** Conceptual Venn diagram showing the ground truth, library, and both derived basis sets. **B.** Overlap of the derived basis sets compared to the ground truth for the healthy spectra (top) and tumor spectra (bottom). From left to right, columns show the number of false positives, the number of false negatives, and the Sorensen-Dice coefficient (SDC) reported as a percentage. Within each panel, the left half shows the 100 single-spectrum results, and the right half shows the sole result of the group-wise optimization. Stopping conditions are color-coded: Max-AIC (green) and zero-Amp (orange). Below each bar, the mean value is reported.

For group-wise optimization of healthy spectra, both stopping conditions converged to the same basis set composition. For the tumor spectra, the max-AIC stopping condition included 4 additional metabolites, 3 of which were not present in the simulation. Across both datasets and both stopping conditions, the group-level optimization achieved higher SDC than the single-spectrum optimization. The ground-truth overlap increased by 6% for the max-AIC (from 86% to 92%) and 8% for the zero-amplitude stopping conditions (from 84% to 92%) in the healthy spectra, and increased by 1% and 5%, respectively, for the tumor spectra.

When linewidth and noise were increased, the algorithm suggested sparser basis set compositions, as the model was able to utilize the baseline and linebroadening to adequately (according to the AIC) model the broad, noisy spectra with fewer components. In contrast to the high-quality datasets, the single-spectrum optimization was more robust than the group-wise optimization, maintaining >73% overlap with the ground truth basis set for even the worst-quality data (zero-amplitude, single-spectrum; Figure 5).

**Figure 5.**
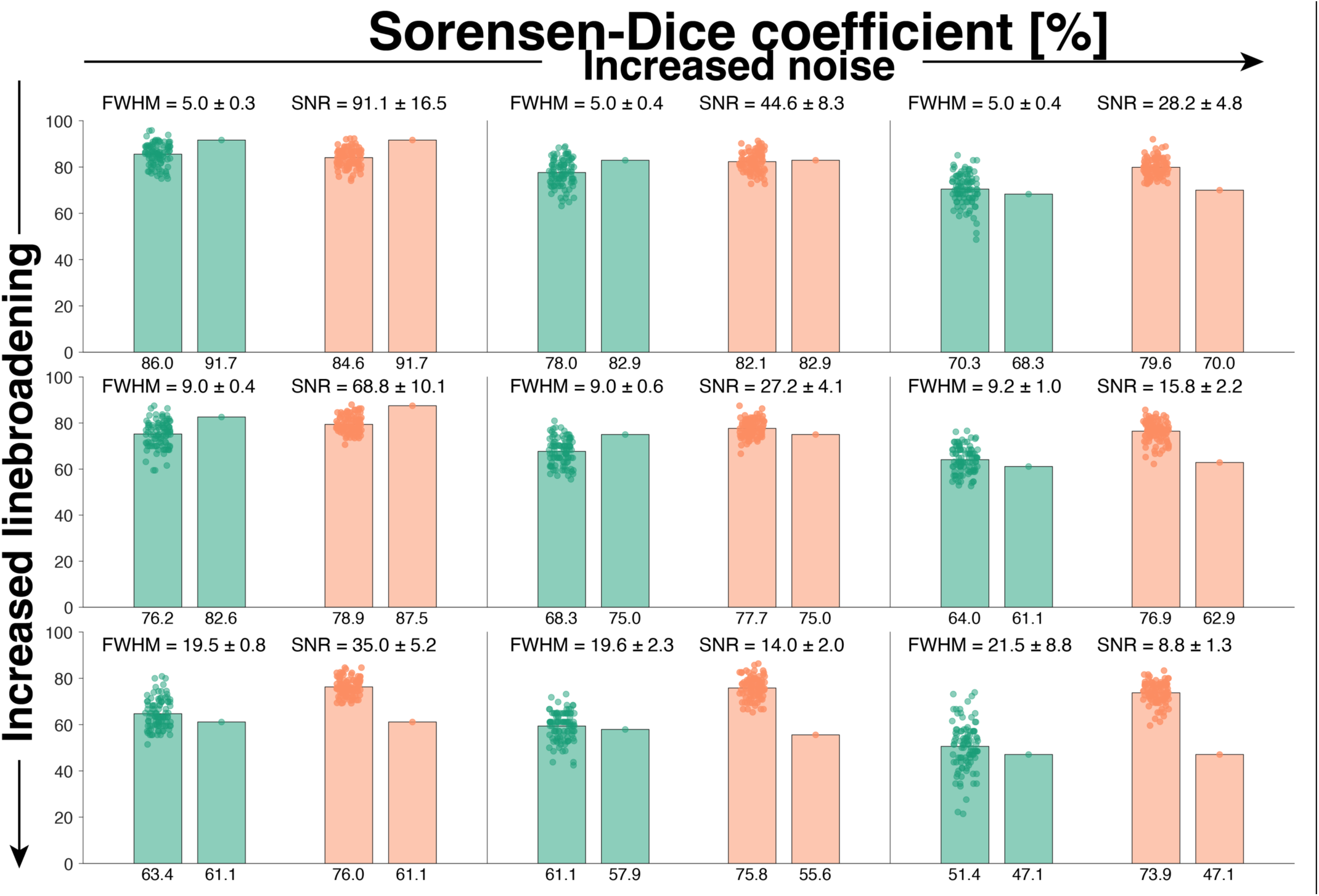
Sorensen-Dice coefficient (SDC; percentage) for each of the 9 data-quality scenarios. Bars are colored by stopping condition—Max-AIC (green) and zero-Amp (orange)—with the single-spectrum optimization on the left (multiple data points) and group-wise optimization on the right (single data point). Labels on the x-axes indicate the median of the distributions.

### 3.3. Inclusion of 2HG and Cystat

For the single-spectrum optimization, the inclusion or omission of Cystat and 2HG depended on the stopping condition used, with neither providing perfect results. The zero-amplitude condition included the oncometabolites in 97% of the tumor spectra (n_Cystat_ = 100; n_2HG_ = 94) but with many false positives for these same metabolites in healthy spectra (n_Cystat_ = 54; n_2HG_ = 42). The situation is somewhat reversed for the max-AIC stopping condition. It does better at omitting Cystat and 2HG from healthy spectra (n_Cystat_ = 5; n_2HG_ = 10) but includes them for fewer (92%) tumor spectra than zero-amplitude (n_Cystat_ = 95; n_2HG_ = 89). Notably, all of the excluded “false negatives” had very low ground-truth amplitude, suggesting a (fairly low) amplitude threshold above which the algorithm almost always correctly included Cystat and 2HG (Figure 6B).

**Figure 6.**
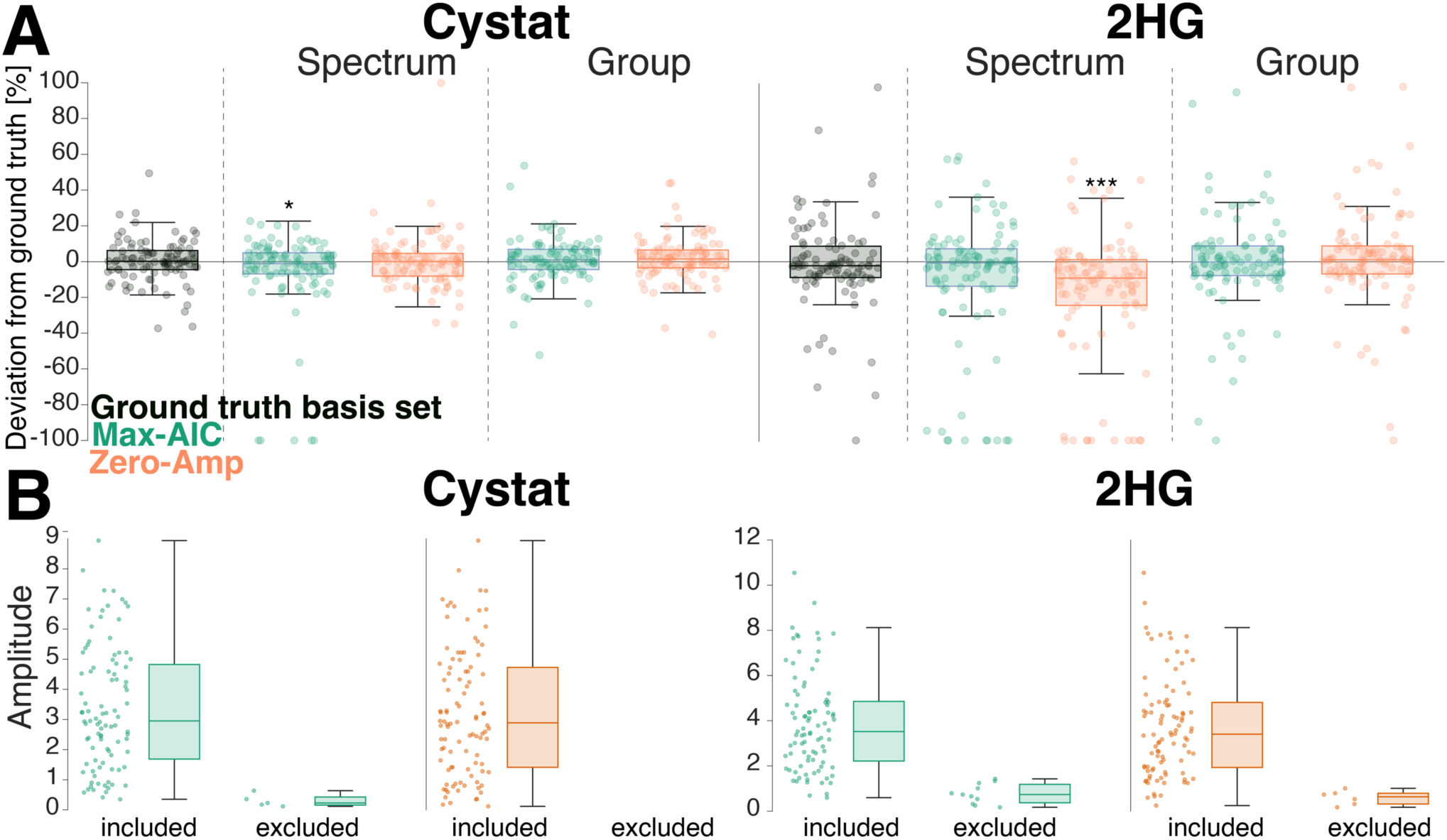
**A.** Boxplots of the percentage deviation of amplitude estimates from the simulated ground truth for Cystat and 2HG in the tumor spectra. The results using the ground truth basis set (i.e. including only the metabolites present in the simulation) are shown in black. The two stopping conditions—max-AIC (green) and zero-Amp (orange)—are shown in each panel, for the single-spectrum optimization (left) and group-level optimization (right). Significant deviations from the estimates of the ground-truth basis set are marked with asterisks: “*” (0.005 ≤ *p* < 0.05), “**” (0.0005 ≤ *p* < 0.005), and “***” (*p* < 0.0005). Note that an estimated amplitude of zero is assumed for single-spectrum-optimized basis sets which omitted that particular metabolite. This resulted in a cluster of data points at –100%. However, 4/10 of these points for the single-spectrum, zero-amplitude basis sets were true zero-amplitude estimates of 2HG when it was present in the derived basis set. **B**. Boxplots showing the ground-truth amplitude distributions of Cystat (left panel) and 2HG (right panel) for the synthetic tumor spectra. Boxplots are separated by the respective inclusion/exclusion (left/right) of the Cystat or 2HG in the derived basis sets derived for single-spectrum optimization with the zero-amplitude (orange) and max-AIC (green) stopping conditions.

For group-level optimization, the results are encouraging and definitive. Cystat and 2HG were correctly omitted from the basis set of the healthy spectra and correctly included in the tumor spectra for both stopping conditions.

### 3.4. Effect on metabolite amplitudes

Metabolite amplitude errors depended on the ground truth amplitude, with larger relative error for low-concentration metabolites. Figure 7 shows the percentage deviation of metabolite amplitude estimates from the ground truth for 5 metabolite measures commonly reported in the MRS literature: tNAA (NAA + NAAG), tCr (Cr + PCr), tCho (GPC + PCho), mI, and Glx (Glu + Gln).

**Figure 7.**
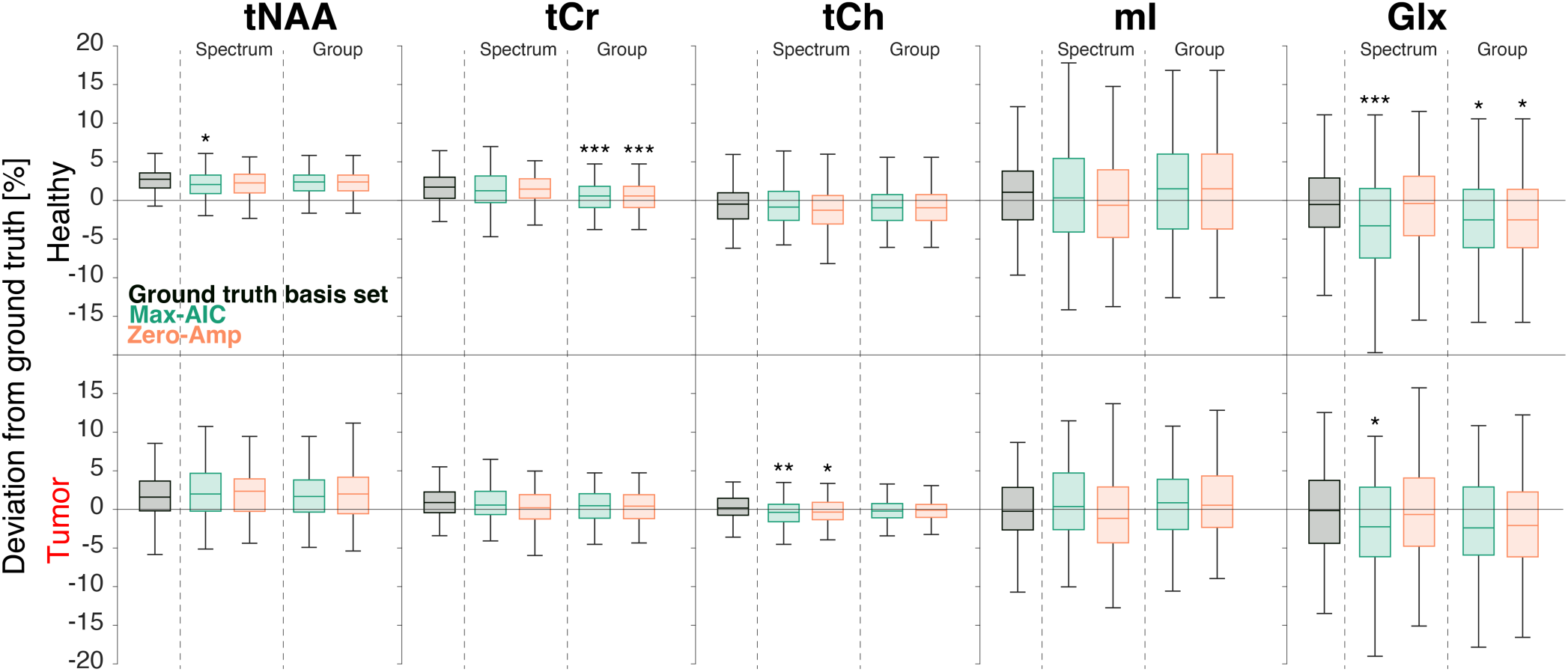
Boxplots of the percentage deviation of amplitude estimates from the simulated ground truth for 5 commonly reported metabolites (tNAA, tCr, tCho, mI, and Glx). Results for the healthy spectra are shown in the top row and tumor spectra in the bottom. The results using the ground truth basis set (i.e. including only the metabolites present in the simulation) are shown in black. The two stopping conditions—max-AIC (black) and zero-Amp (orange)—are shown in each panel, for the single-spectrum optimization (left) and group-level optimization (right). Significant deviations from the estimates derived using the ground-truth basis set are marked with asterisks: “*” (0.005 ≤ *p* < 0.05), “**” (0.0005 ≤ *p* < 0.005), and “***” (*p* < 0.0005). Note that the plot limits exclude Glx outliers for comparability.

Absolute amplitude errors were typically below 15% for these 5 metabolites, with some negative Glx outliers as large as –32%, namely for max-AIC-derived basis sets. For the major singlet resonances (tNAA, tCr, and tCho), absolute amplitude errors were < 17% overall, and ≤ 11 % for the healthy-appearing spectra, specifically.

3 out of 4 algorithm variations achieved the same average mean amplitude error (3.2%). The exception—max-AIC; single-spectrum optimization—predicted somewhat sparser basis sets and exhibited a larger mean amplitude error (3.5%). The best-performing optimization method (as measured by SDC; group-wise & zero-amplitude) only deviated from the ground truth amplitude estimates in healthy spectra for Glx (mean difference = 2.1%, *p* = 0.0071) and tCr (mean difference = 1.2%, *p* < 0.0005).

When the spectral quality of the synthetic data was reduced, amplitude estimates were robust to noise, but we observed significant underestimation of metabolite amplitudes in datasets with increased linewidth (tNAA, tCr, tCho, and mI), even when the correct basis set was used (**Figure S10**). In these datasets, the MRS model increasingly assigned signal to the baseline and other broad basis functions, particularly lipids and MMs. Interestingly, while the sparse algorithm-derived basis sets exhibited broader metabolite amplitude error distributions (lower precision), the median errors were generally closer to zero (improved accuracy). This suggests that despite the limitations, there may be a precision-accuracy trade-off for using this approach in low-quality spectra.

The amplitude results for Cystat and 2HG are shown in Figure 6A for the tumor spectra. All conditions achieved median ground-truth errors of less than 10% for both 2HG and Cystat, with small (but significant) deviations from the ground-truth basis set estimates. Both group-based optimizations estimated 2HG and Cystat without significant differences to the ground-truth basis set.

When we compared the amplitude error distributions of the basis set derived by our algorithm to the fixed expert-recommended compositions, our method performed favorably (Figure 8). Amplitude error distributions varied greatly across the expert-recommended basis sets, with some exhibiting large systematic offsets. For example, basis set 3 omitted NAAG and GSH, both present in the simulation, and so the model erroneously assigns signal to NAA and Glx to compensate. In almost all cases, the amplitudes predicted by the algorithm-derived basis set better agreed with those of the known ground truth basis set.

**Figure 8.**
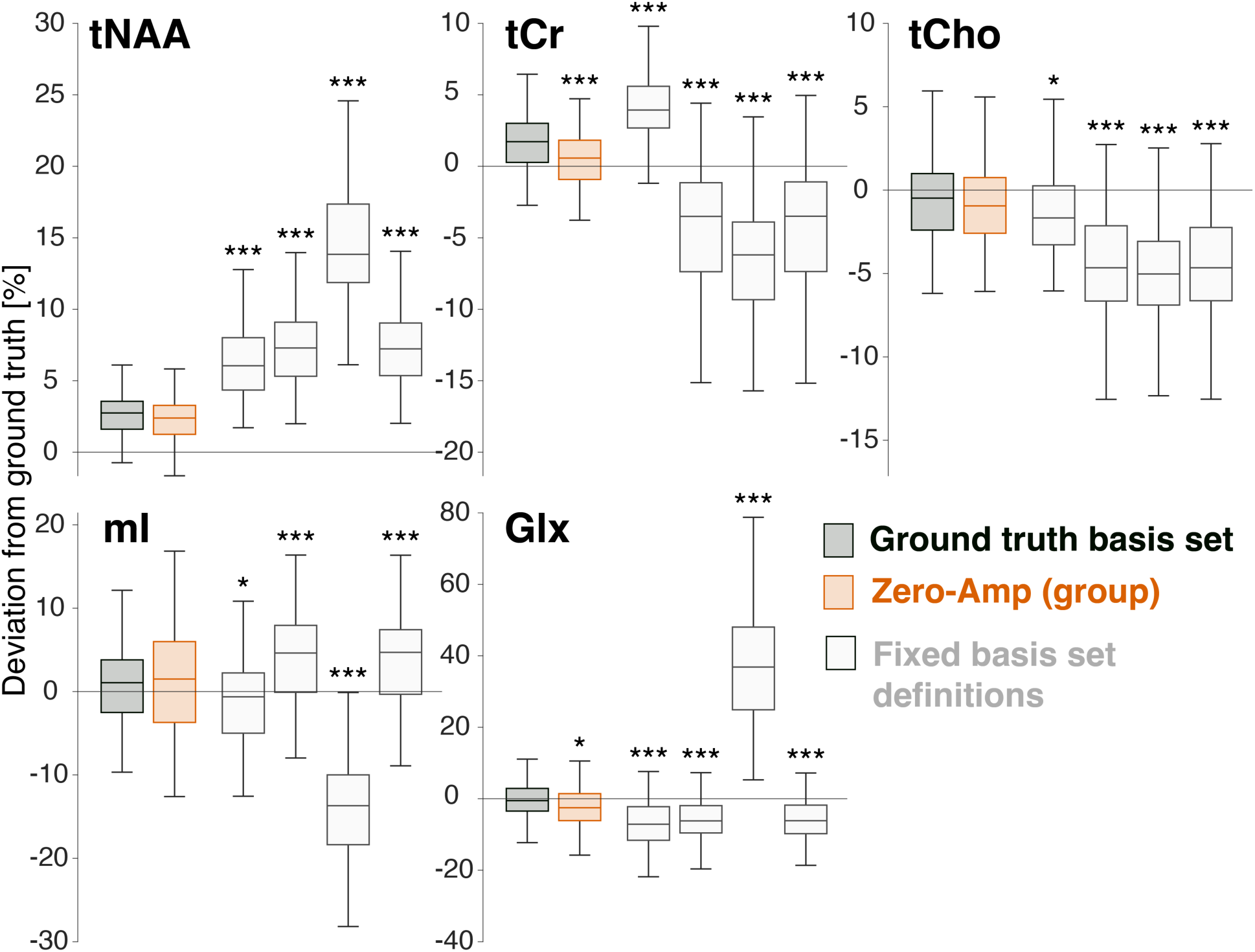
Boxplots showing the distribution of metabolite amplitude errors derived for each basis set: the correct ground truth set (black), the groupwise zero-amplitude algorithm-derived basis set (orange), and the four expert basis set definitions (light grey). Significant deviations from the ground truth basis set (paired t-test) are labeled with asterisks.

### 3.5. In-vivo data

Both single-spectrum and group-based flavors of our algorithm selected most metabolites that are commonly encountered in in-vivo basis sets (Figure 9**; Figure S11**). Notable omissions from the group-wise-optimized basis set were MM09, MM12, and Lip20, likely because they are sufficiently modeled by similar-appearing broad signals (Lip09, Lip13, MM20). Similarly, EtOH and Cit were surprisingly included, likely because they can resemble aspartyl signals around 2.5 ppm (Cit) and, if sufficiently broadened, MM/Lip signals around 1.2 ppm (EtOH); however, EtOH was included in the final round of the (more aggressive) max-AIC stopping condition.

**Figure 9.**
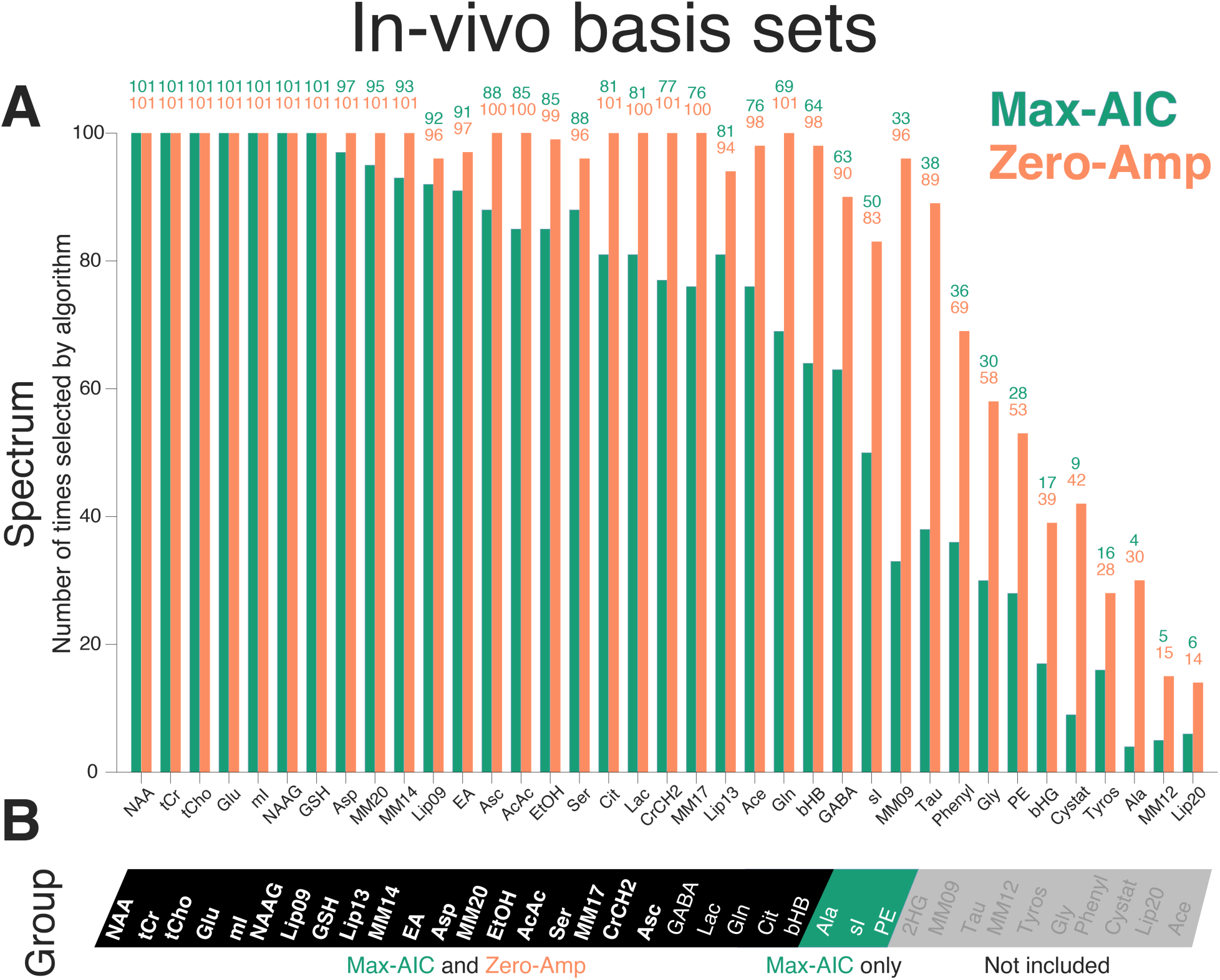
Basis set composition derived from short-TE in-vivo data. **A.** Bar chart showing the number of times each metabolite was selected for inclusion in the single-spectrum-optimized basis sets for the in-vivo data. Colors represent the two stopping conditions. Metabolites are listed in descending order according to the total number of inclusions across both stopping conditions, i.e., more likely included (left), less likely (right). **B.** A representation of the metabolites selected from the same data as above using group-wise optimization. The black panel indicates metabolites included in both stopping conditions, the green panel shows metabolites included only in the max-AIC stopping condition, and the light grey panel indicates metabolites excluded by both stopping conditions.

## 4. Discussion

A key challenge for in-vivo MRS is that a ‘true’ model of the spectrum cannot be known, since the processes that shape it—the interplay of concentrations, microstructure, microscopic particle motion, macroscopic subject motion, temperature, pH, etc.—are too complex. All conventional ^1^H-MRS modeling methods therefore focus on the most relevant macroscopic quantities—i.e., signal amplitudes for key metabolites—and simplify other aspects. This epistemic uncertainty has provided a fertile environment in which a multitude of model functions, algorithms, software tools, and fitting strategies have grown over the last thirty years. Unsurprisingly, the different models and strategies do not agree very well with each other^28–30^, contributing to the poor reproducibility and comparability of metabolite estimates in the MRS literature^31^.

“If we have several options to model our data, which should we pick?” is a common question across the sciences. If there are multiple suitable ways to model in-vivo MRS data, we posit that modeling procedures should not only traverse the parameter space *within* a single model definition (via least-squares optimization) but also search *across* model spaces, i.e., explore the breadth of reasonably admissible models. Quantitative model selection methods provide a formal, rigorous framework to do this—exemplified by the ABfit algorithm, which selects from many admissible baseline spline models, affording just enough model flexibility to approximate the data, but not more^17^.

The composition of the basis set is another prime example of the range of reasonably admissible models, having been at the sole discretion of individual researchers since the inception of LCM for in-vivo MRS. The surprisingly limited number of studies that have investigated this modeling decision highlights the potential for operator bias, with changes to the basis set composition substantially affecting metabolite estimates^4–10^. In this proof-of-concept study, we therefore demonstrate the use of model selection for determining basis set composition directly from the spectra.

Across all procedural variations, the algorithm derived reasonable basis sets and consistently produced fits with flat residuals. High-concentration metabolites were correctly included in the algorithm-derived basis sets, without exception. The algorithm’s ability to correctly identify lower-concentration metabolites depended on the precise procedural variation and quality of the data. We found that group-level basis set estimation with the ‘zero-amplitude’ stopping condition minimized the number of false positives and negatives, correctly recognized 2HG and Cystat in the tumor group, and provided metabolite estimates close to those estimated with the ground-truth basis set, at least in the high-quality data. This is an encouraging starting point, although further investigation is required.

### 4.1. Choice of Stopping Condition

We initially designed the algorithm to select the basis set that maximized the BIC score— certainly the most intuitive choice, as it is purportedly the optimal balance of model complexity and fidelity, and the BIC exhibited the largest dynamic range of scores (**Figure S3**). However, the BIC scores peaked too early, including too few metabolites in the basis set (**Figure S12**). The BIC penalized additional metabolites too heavily and the least-squares model was flexible enough to mimic the signal of metabolites (particularly J-coupled, low-concentration ones) through some combination of baseline bumps, lineshape distortions, and overlapping metabolites already present in the basis set. This—according to raw BIC scores—is preferable to including another basis function, which adds model parameters. Switching to the AIC mitigated this by reducing the penalty on including additional metabolites.

Our other (non-IC) stopping condition added further metabolites (beyond the IC maxima) until all remaining basis functions are fit with zero amplitude. This “zero-amplitude” approach agreed well with the max-AIC stopping condition in the group-wise optimizations—even improving things in the tumor spectra—but resulted in more false positives when the optimization was performed on a per-spectrum basis.

### 4.2. Group-wise or single-spectrum

Group-wise optimization improved the identification of lower-concentration metabolites. The aggregation of AIC scores across many spectra reduced the impact of noise and individual amplitude variations, allowing the algorithm to better prioritize signals that are inconsistently modeled in a single spectrum. However, single-spectrum optimization performed better for low-SNR and high-linewidth spectra, suggesting this benefit of group-wise optimization is limited to consistently high-quality datasets (Figure 5 & **Figures S5–S10**). Group-wise optimization also removes our ability to adapt to individual outlier spectra that may be present in heterogeneous in-vivo samples. To overcome this limitation, ‘soft inclusion’ methods (machine learning or traditional regularization strategies) may harness the probability of the presence of metabolites from the full group and then include specific metabolites in single-spectrum fits if sufficiently justified by the data, rather than ‘hard exclusion’ of a metabolite (see Section 4.5 *Perspectives*).

Future work could incorporate additional information into the decision-making process to tackle issues of metabolite overlap. These may include Cramér-Rao lower bounds or metabolite amplitude correlation matrices derived from the Fisher information matrix, or it may be beneficial to modify the IC itself, changing the relative weighting of the terms representing the model residual and number of parameters. Optimization of this balance might improve the max-IC stopping condition, but further work is needed to explore this.

The correct inclusion of 2HG and Cystat in the tumor cohort (and simultaneous exclusion in the healthy cohort) illustrates that the selection process might even deem two different models to be most appropriate if two cohorts exhibit markedly different spectral characteristics. This is, in general, of course, desirable! For example, healthy tissue does not produce detectable levels of 2HG^32(p2)^, and model selection for a set of healthy spectra *should* select a model that does not contain a 2HG basis function because fewer degrees of freedom will reduce the variance in estimating signals like Glu/Gln that overlap with 2HG. However, the interaction between these signals is acquisition-dependent, and so the inclusion of 2HG—even when it is present—will depend on the degree to which we can independently model its signal.

### 4.3. The impact of linebroadening and noise

In low-quality spectra where LCM was challenging, our algorithm was less effective in correctly identifying the ground truth basis set. As linewidth and noise increased, the algorithm-derived AIC curves tended to follow flatter trajectories that peaked earlier (**Figures S6–S7**). The model was able to reasonably approximate the broader spectra using fewer basis functions in combination with the baseline, and noise obfuscated the contributions of lower-concentration signals. The algorithm-derived basis sets were therefore sparser for poor-quality data and exhibited an increased number of false negatives (**Figures S8–S9**).

For the broadest spectra, metabolite amplitudes were estimated with larger errors. Metabolite signal was erroneously assigned to broad lipid/MM basis functions and the baseline. However, this was true for algorithm-derived basis sets, models using the full library set, and even when the correct ground-truth basis set was used (**Figure S10**). This suggests that the amplitude results were characteristic of the model implementation itself, rather than the algorithm. Compared to the ground-truth basis set, the algorithm-derived basis sets estimated metabolites with lower precision (larger dispersion of errors) but often with higher accuracy (distributions were closer to zero). Despite these encouraging signs—and avenues for future development—we would not currently recommend this approach for anything other than high-quality data. It should, however, be noted that current practice encounters the same problems—researchers tend to include fewer basis functions when analyzing poor-quality data, but the exact choice remains at their individual discretion, and no formal guidance or evidence exists.

### 4.4. Limitations

Our synthetic data omitted some common characteristics of in-vivo MRS data. For example, we did not introduce phase alterations, frequency shifts, correlated noise, or baseline components. Broad resonances underlying the metabolite signals, in particular, significantly impact the accuracy of MRS modeling^8,29^. Baseline modeling would be further complicated by the frequent omission of MM and lipid basis functions. The algorithm tended to capture some of these— admittedly small—signals using the baseline, reducing the number of model parameters, but compromising the validity of the baseline model. This inability of the model to correctly characterize the lipids and MMs also undermined the analysis of the heavily-linebroadened spectra. Future work could mitigate this by augmenting the relatively simple model definition used in this study. For example, we could introduce soft constraints on relative metabolite and MM amplitudes, include some algorithm penalty for baseline amplitudes, introduce additional modeling steps to improve global lineshape characterization, or use experimentally acquired MMs, as recommended by expert consensus^33^ and our previous investigations^29,34^.

Furthermore, careful considerations need to be made for signal artifacts. Out-of-voxel “ghost” echoes introduce unpredictable signals that cannot be accurately captured by the basis set. Large, broad signals might also be introduced by extraneous lipids or residual water signals, which get incorporated into the model’s spline baseline or existing lipid basis functions. Our approach could misattribute metabolite basis functions to these signals, so specific measures would need to be implemented to avoid this, preferably during data acquisition or, if need be, with specific post-processing techniques^35,36^ before applying our algorithm.

Lastly, the repeated modeling required for this method is far more computationally expensive than the traditional single-basis-set approach. For a library set of size *N*, our “greedy” algorithm requires 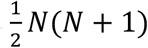 model calls to complete the full IC curves, like those shown in Figure 2. While this is still significantly faster than an exhaustive search of all possible basis sets (requiring 2*^N^* model calls—a factor of 1.4 × 10^’^ slower for N = 40), there is also a possibility of not establishing the global IC optimum with our approach. To improve the computational efficiency of our method, one approach could be to use this algorithm to refine some expert-defined basis set, adding (or subtracting) case-specific metabolites like 2HG or Cystat.

### 4.5. Perspectives

Model selection can, in principle, be applied to any aspect of modeling. Information criteria may be useful to determine optimal lineshape representations, e.g., deciding whether simple Voigtian parametrization suffices, or whether a more generalized convolution kernel (with more model parameters) is required to adequately fit lineshape irregularities resulting from B_0_ inhomogeneities. They may also help decide whether metabolite-specific frequency shifts are warranted by the data or whether a single, global, shift parameter suffices.

Parameter regularization may provide an alternative to model selection. This usually involves additional terms in the model expression that penalize the deviation of model parameters from expectation values or impose constraints on, e.g., smoothness. This often-overlooked injection of prior knowledge is widely considered to be the “secret ingredient” of LCModel and is not well studied. Future work should, therefore, explore whether this approach can complement model selection. For example, instead of excluding certain metabolites like 2HG altogether, it might be wiser to simply set their amplitude expectation values to zero. This would incentivize the model to only assign signal to 2HG when it is strongly warranted by the data, akin to approaches such as “LASSO”^37^. Of course, choosing the nature and strength of the regularization terms is, yet again, a form of operator bias.

Finally, it is important to note the influence of the model, itself. A modification to—e.g. regularization of the metabolite amplitudes or baseline—will inevitably alter the IC scores, and, eventually, the derived basis set. Indeed, the best solution to the epistemic uncertainty problem may be to not even try to select a *single* model but to synthesize the evidence from *multiple* models. So-called multiverse analyses^38–42^ have been pioneered to integrate multiple statistical models in psychology and have recently been applied to resting-state fMRI analysis. It has further been proposed to weigh different models by their Akaike information criterion scores^43–45^ to emphasize the evidence from more parsimonious models, conceptually similar to CRLB weighting to improve statistical inference across multiple spectra^46^.

## Conclusion

Data-driven determination of the basis set composition is feasible with single-spectrum and group-level optimization. With refinement, this method could provide a valuable data-driven way to derive or refine basis sets, reduce the operator bias of MRS analyses, enhance the objectivity of quantitative analyses, and increase the clinical viability of MRS.

## Consent for publication

All authors consent to the publication of this study.

## Data availability

The synthetic data, the means to generate it, the algorithm, and all supplementary code are freely available online: DOI 10.17605/OSF.IO/P2USJ.

---

**Table S1.**
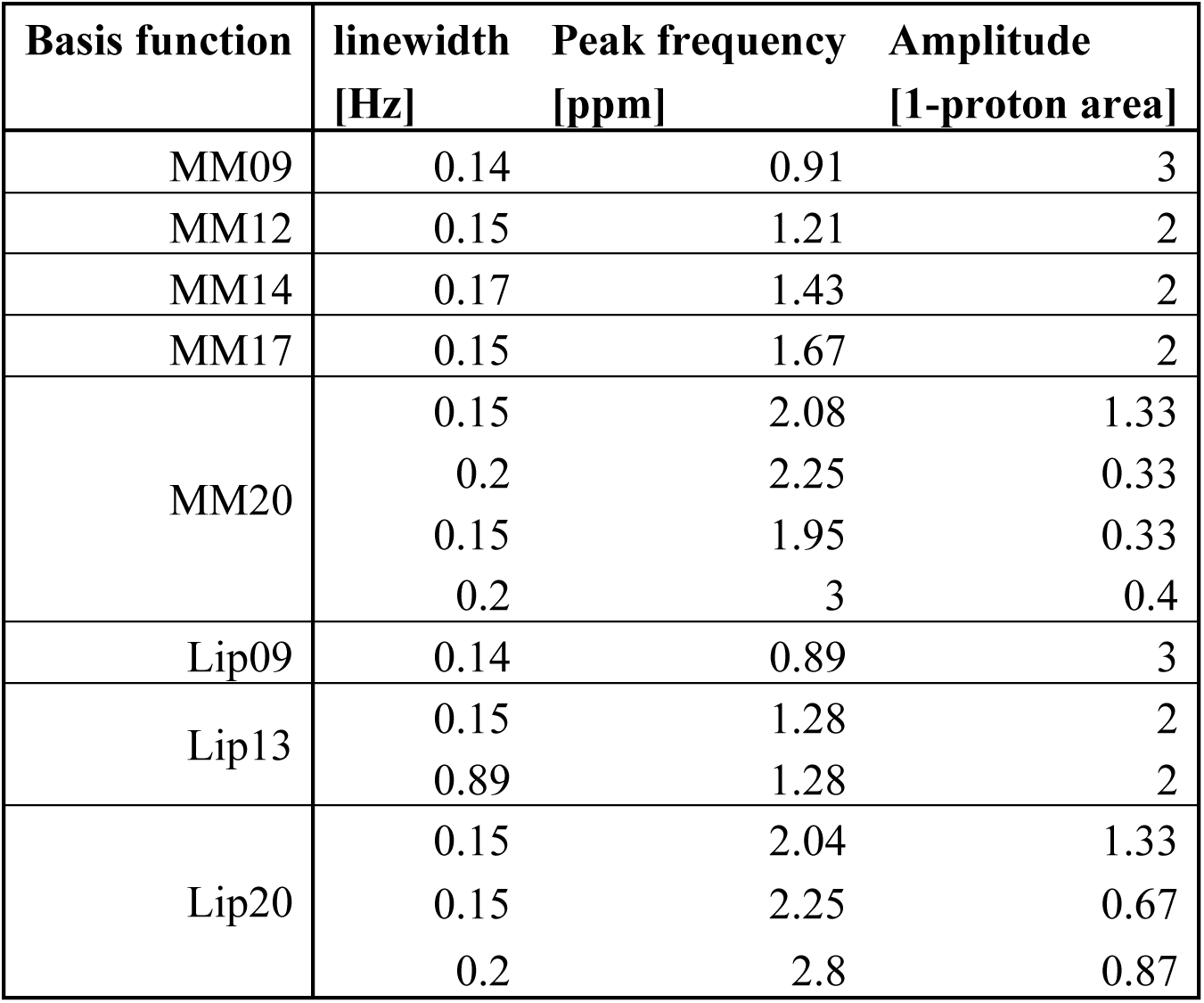
The width, center frequency, and relative amplitude of the parameterized lipid and MM resonances we included in the basis set. Specific values were based on the LCModel parameterizations of these signals.

**Figure S1.**
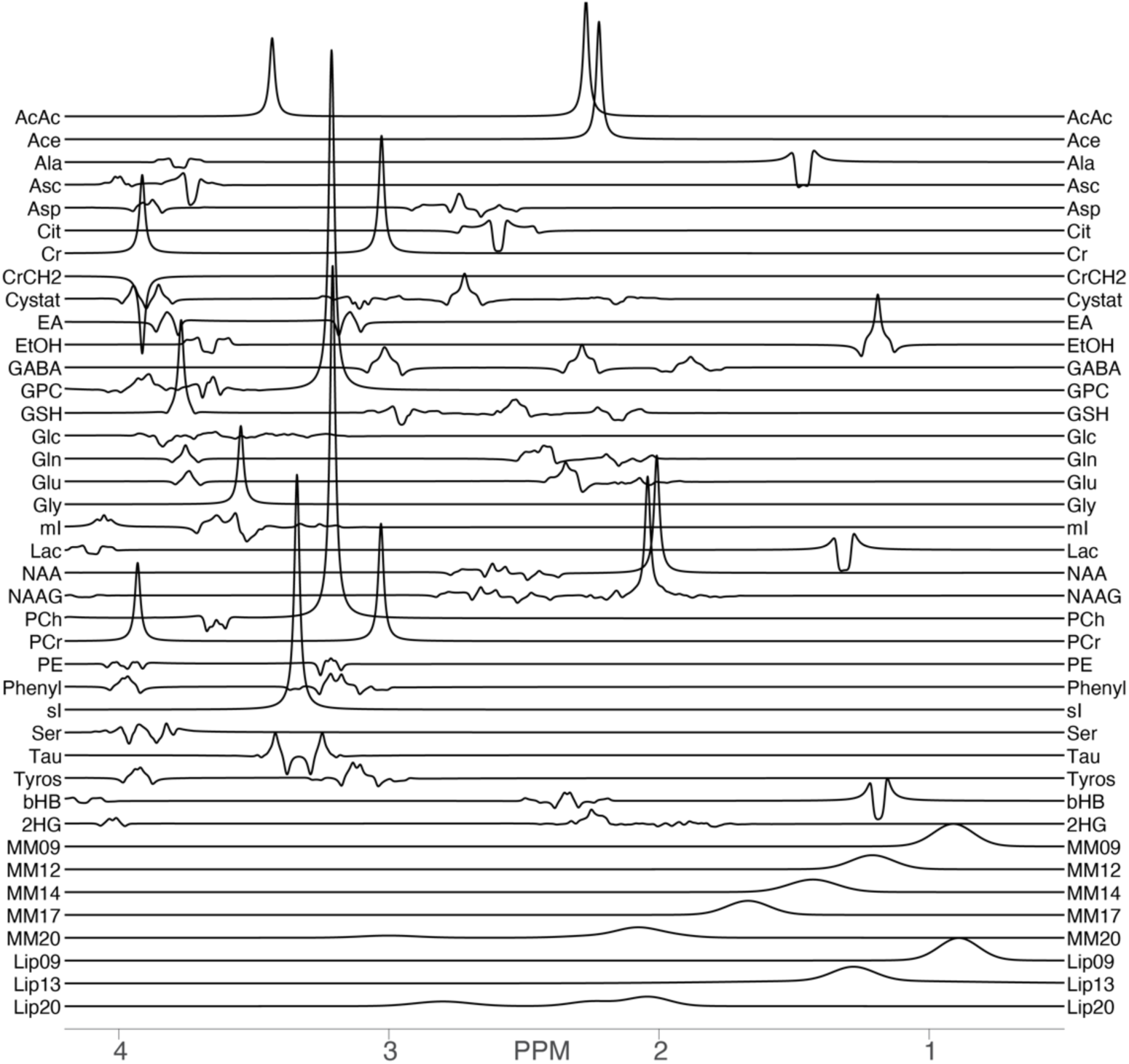
The full “library” basis set of metabolite basis functions with 1.3 Hz Lorentzian linebroadening.

**Table S2.**
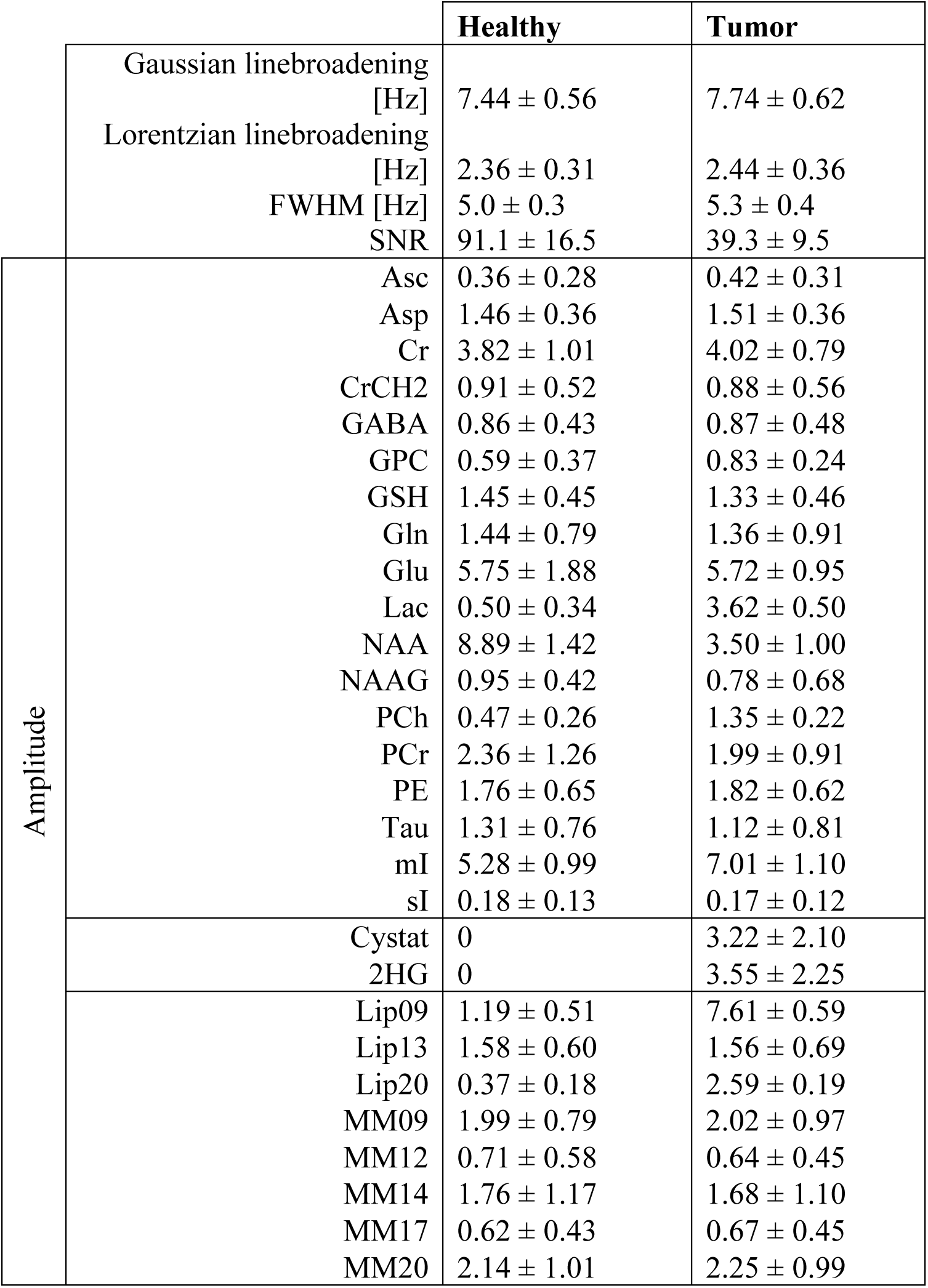
Ground truth simulation parameters (mean ± standard deviation) used to derive the healthy-appearing spectra (left column) and low-grade glioma, “tumor” spectra (right column). The top three rows show the Gaussian and Lorentzian linebroadening terms and the SNR, with subsequent rows showing the metabolite-specific amplitudes.

**Figure S2.**
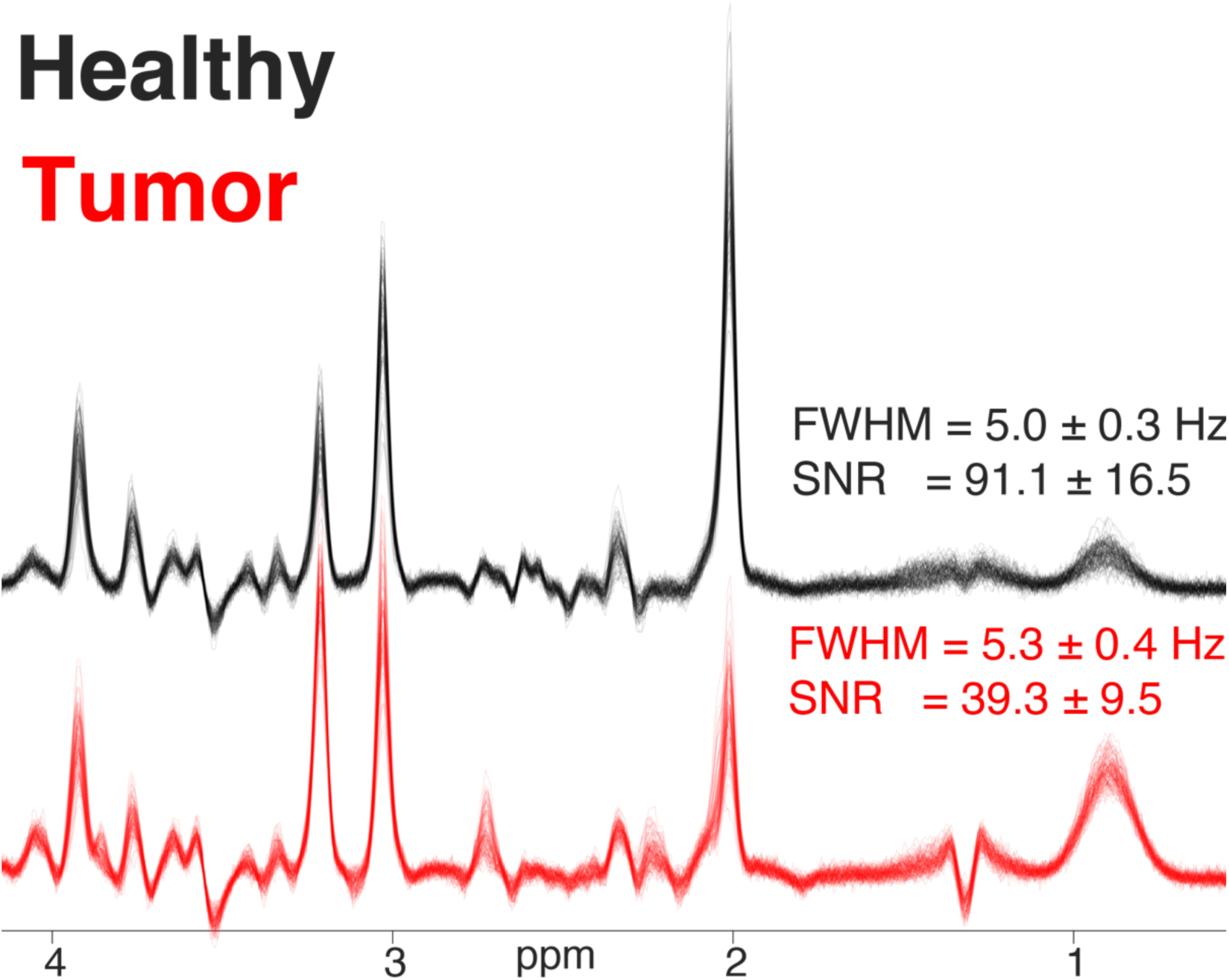
The synthetic dataset used to test the algorithm. Healthy-appearing spectra (black) are plotted alongside the “tumor” spectra, representing low-grade glioma patients (red).

**Figure S3.**
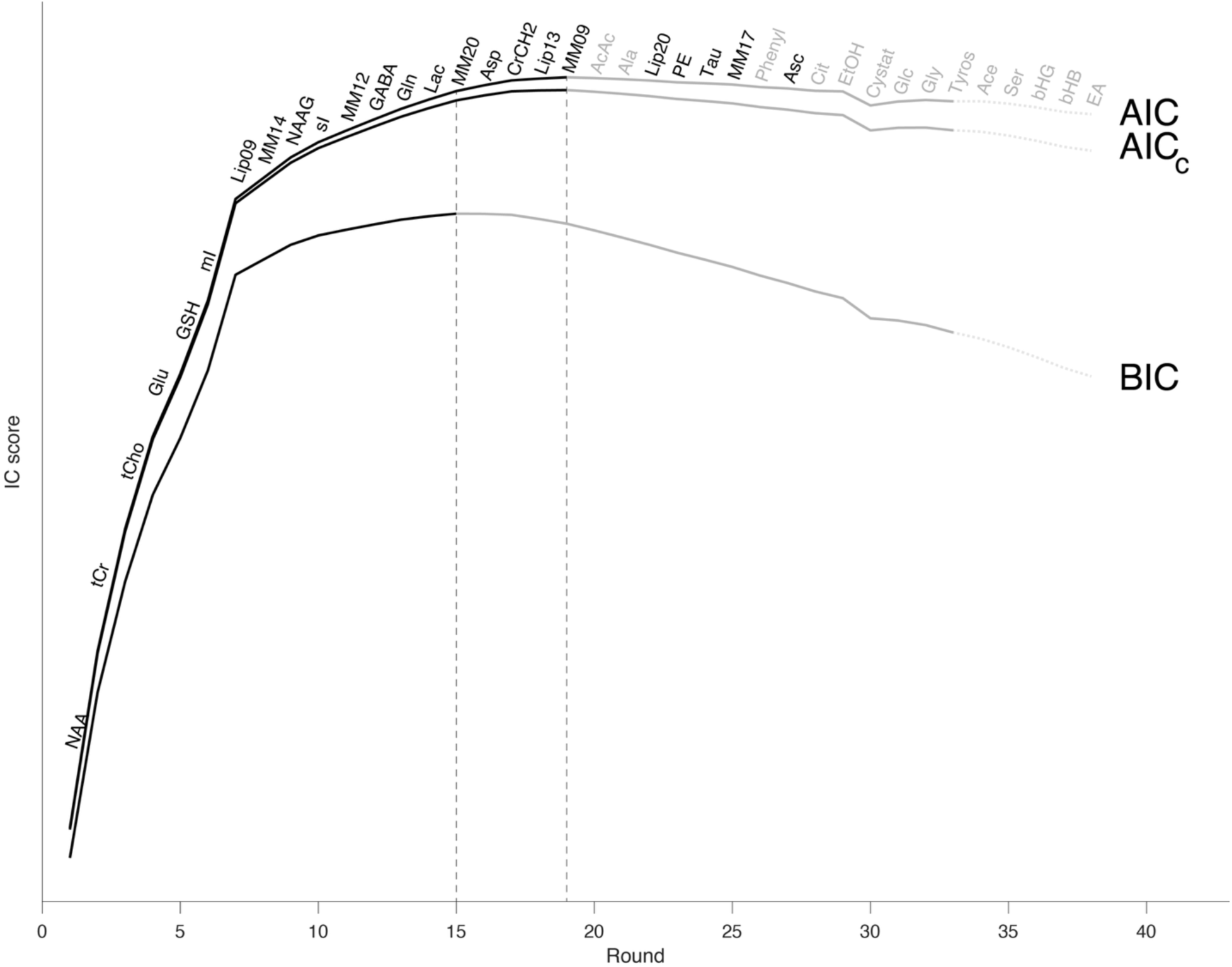
A comparison of IC curves for 3 different IC definitions—the Akaike IC (AIC), corrected AIC (AICc), and Bayesian IC (BIC). There was unanimous agreement between the three metrics regarding the order of metabolites added at each round (text labels), but the BIC exhibited an earlier peak, and therefore, included fewer metabolites in the selected basis set.

**Figure S4.**
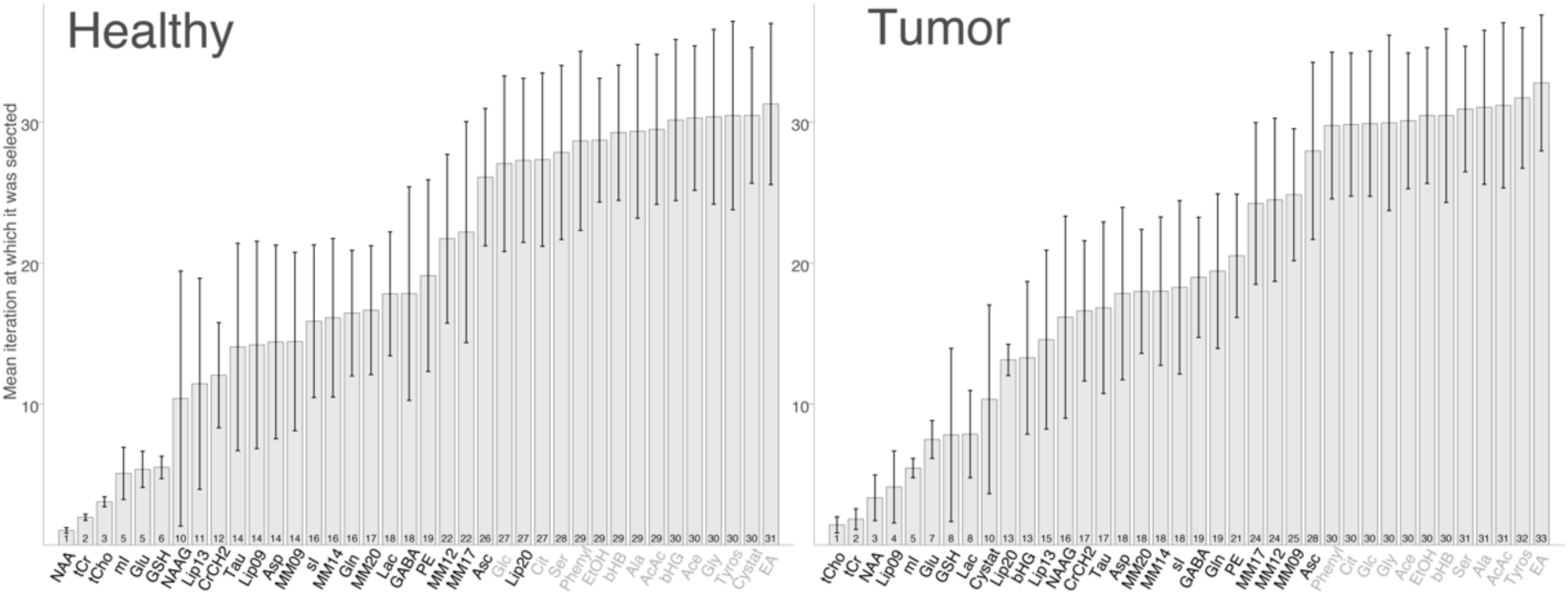
Visualization of the mean (bar) and standard deviation (error bars) of the round at which each basis function was picked. Smaller bars, therefore, indicate a higher prioritization of a given basis function, whilst error bars indicate the inter-spectrum variability of the metabolite’s prioritization. Basis functions are ordered based on their mean. The shade of the metabolite label indicates the presence (black) or absence (grey) of that particular metabolite in the ground truth simulation.

**Table S3.** Lists of the metabolites included in the 4 expert-defined basis sets used for comparison.

| Metabolite | Basis set 1 | Basis set 2 | Basis set 3 | Basis set 4 |
| --- | --- | --- | --- | --- |
| 'Ala' | Yes | Yes | Yes | Yes |
| 'Asc' | Yes | Yes | No | Yes |
| 'Asp' | Yes | Yes | Yes | Yes |
| 'Cr' | Yes | Yes | Yes | Yes |
| 'CrCH2' | Yes | No | No | No |
| 'EtOH' | No | Yes | No | No |
| 'GABA' | Yes | Yes | Yes | Yes |
| 'GPC' | Yes | Yes | Yes | Yes |
| 'GSH' | Yes | Yes | No | Yes |
| 'Glc' | No | Yes | No | Yes |
| 'Gln' | Yes | Yes | Yes | Yes |
| 'Glu' | Yes | Yes | Yes | Yes |
| 'Gly' | Yes | Yes | No | Yes |
| 'Lac' | Yes | Yes | No | Yes |
| 'NAA' | Yes | Yes | Yes | Yes |
| 'NAAG' | Yes | Yes | No | Yes |
| 'PCh' | Yes | Yes | Yes | Yes |
| 'PCr' | Yes | Yes | Yes | Yes |
| 'PE' | No | Yes | No | Yes |
| 'Tau' | Yes | Yes | Yes | Yes |
| 'mI' | Yes | Yes | Yes | Yes |
| 'sI' | Yes | Yes | No | Yes |

## Supplementary Analysis 1: low-quality data

### Methods

The synthetic healthy brain dataset described in the manuscript was augmented to explore the impact of linebroadening and noise. These new data were generated using the FID-A library and the “op_filter” and “op_addNoise” functions, respectively. We generated synthetic data at three additional-linebroadening levels: 0, 5, and 15 Hz, and introduced additional noise at three levels: 1.5e-4, 4e-4, 7e-4 (time-domain standard deviation; note, the non-zero noise in the low-noise category was to compensate for the applied broadening filter). This resulted in 9 sets of 100 synthetic spectra with varying linewidth and SNR—one of which was the healthy dataset in the paper, which represented the highest data quality.

**Figure S5.**
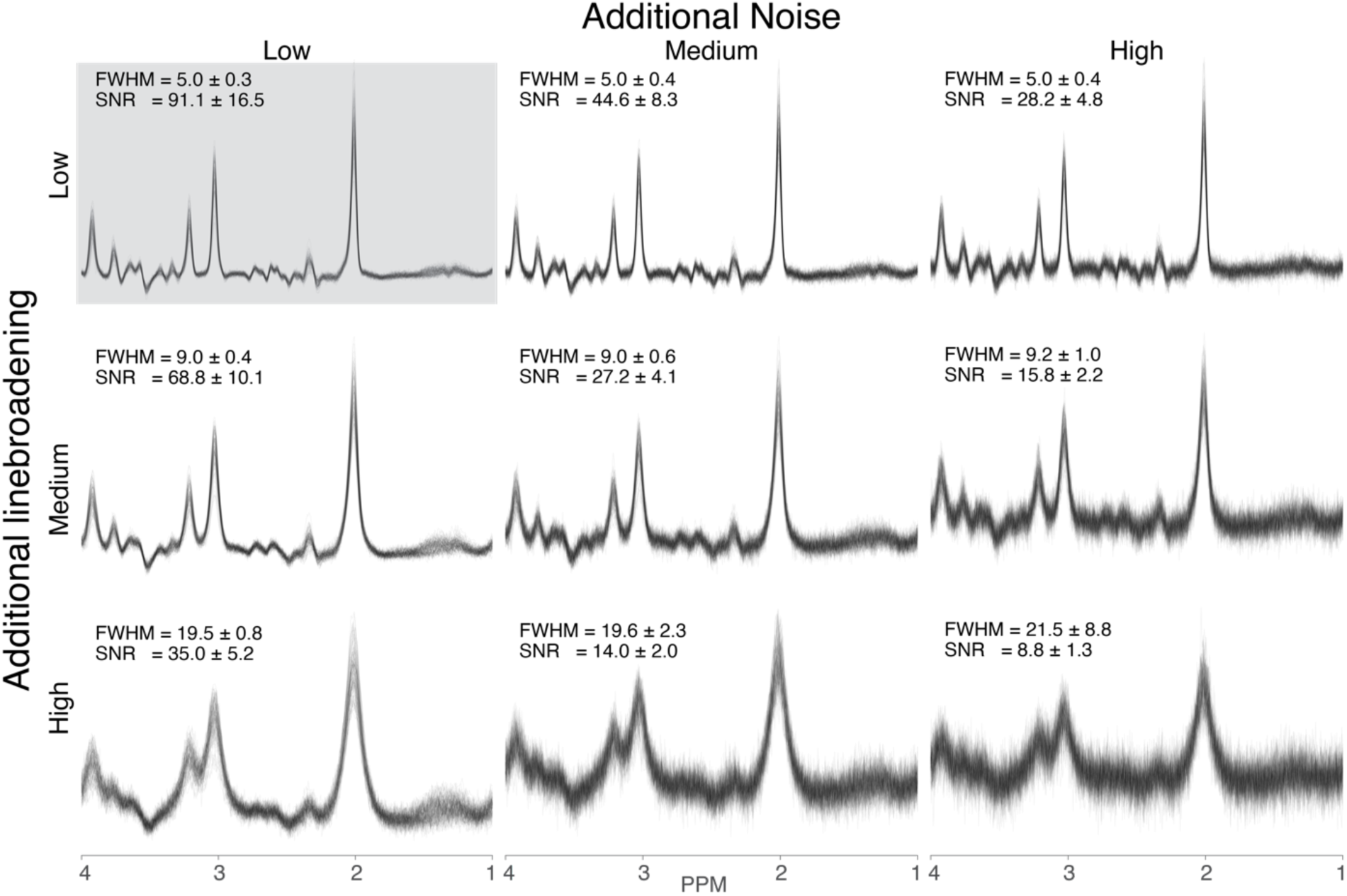
Visualization of the 9 datasets with varied data quality. The top-left panel shows the “healthy brain spectra” from the main study. Lower rows show datasets with increased linewidth, and the right-most columns show datasets with the most additional noise introduced. Labels indicate the distributions of the measured full-width at half maximum (FWHM) and signal-to-noise ratio (SNR) in each case.

We tested the single-spectrum and group-wise variations of our algorithm with both stopping conditions in all 9 datasets. We then examined overlap with the ground-truth basis set using the Sorensen-Dice coefficient and investigated how the amplitude estimates were impacted by basis set changes.

### Results

A qualitative examination of the AIC curves (**Figures S6–7**) reveals a shallower trajectory of the AIC scores as data quality decreases for both single-spectrum and group-level optimizations. The apparent dynamic range of IC scores is restricted as the relative information content of the spectrum is diluted. This results in an earlier maximization of the AIC scores when data quality is low.

**Figure S6.**
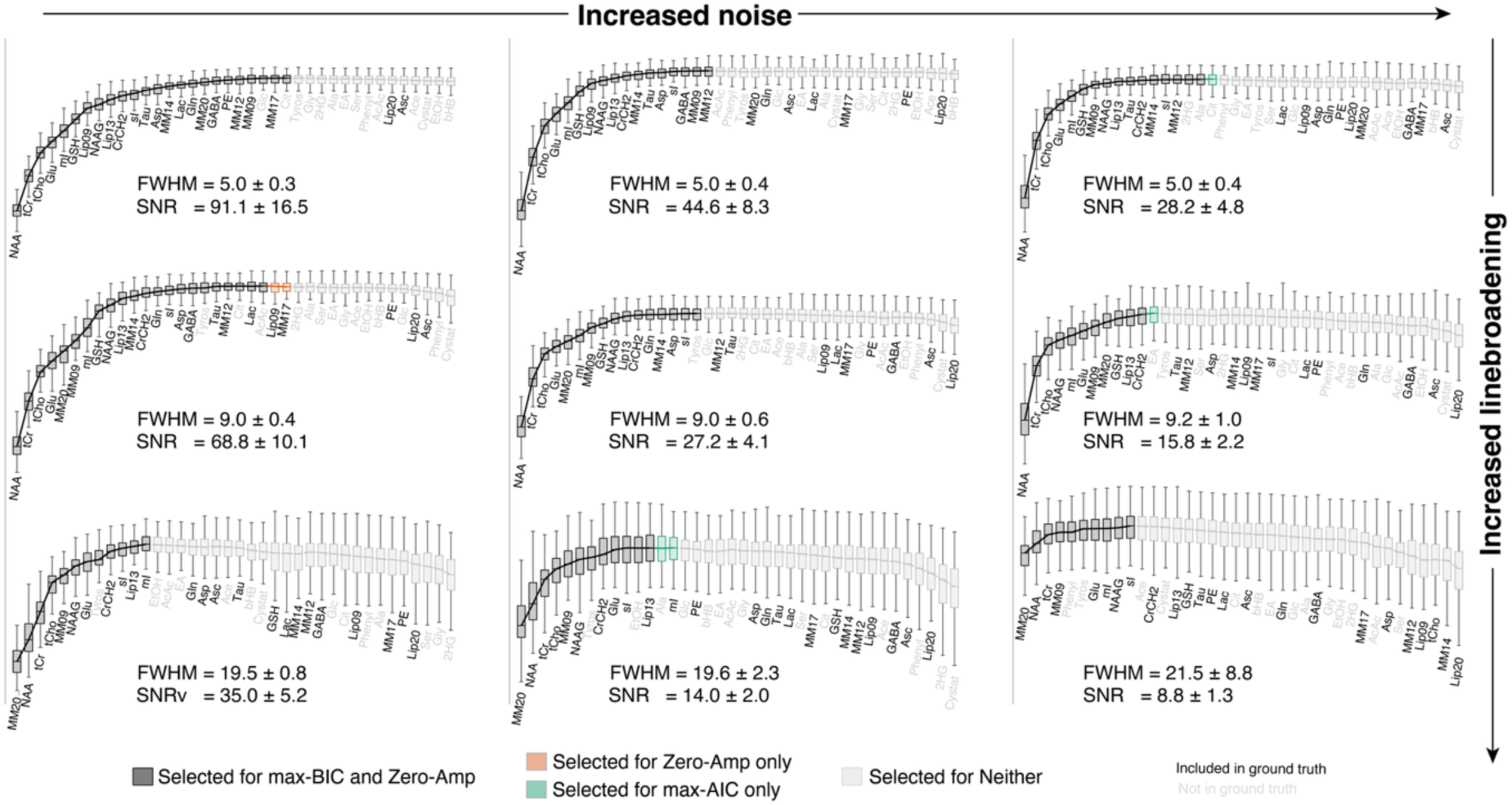
AIC scores as a function of algorithm round for the group-level optimization (common y-scale). Text labels indicate the basis function added at each round, with label color reflecting the presence/absence (black/grey) of that metabolite in the ground-truth simulation. Box plots indicate the AIC distributions across the group at each round. Boxplot color and line style are used to illustrate the extent of the derived basis sets in each stopping condition: included in both stopping conditions (dark box; solid black line), included only in the zero-amplitude condition (orange box; orange line), included in the max-AIC only (green box; green line), or included in neither (light-grey box; light-grey dotted line).

**Figure S7.**
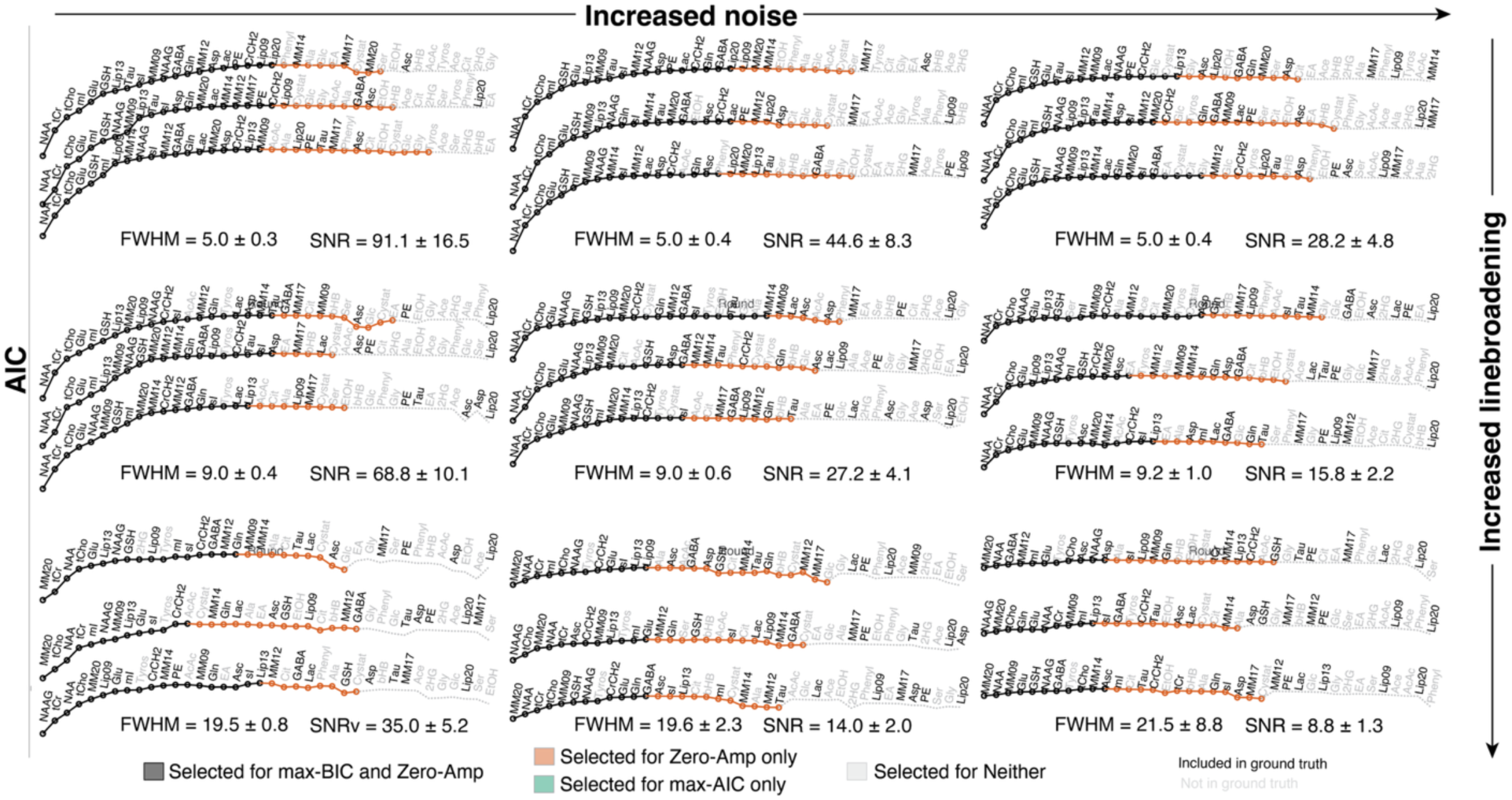
AIC scores as a function of algorithm round for 3 (out of 100) example AIC curves for the single-subject optimizations (common y-scale). Text labels indicate the basis function added at each round, with label color reflecting the presence/absence (black/grey) of that metabolite in the ground-truth simulation. Line style is used to illustrate the extent of the derived basis sets in each stopping condition: included in both stopping conditions (solid black line), included only in the zero-amplitude condition (orange line), or included in neither (light-grey dotted line).

This is precipitated into the quantitative measures of ground truth overlap (**Figures S8–9**). We found that the number of false negatives increases as data quality worsens (**Figure S8**). That is, according to the AIC scores, poorer data are sufficiently described by fewer metabolite basis functions. Conversely, false positives are fairly stable across the differing data quality scenarios (***Supplementary Figure 9***). The net result is an overall decrease in the SDC (Figure 5).

**Figure S8.**
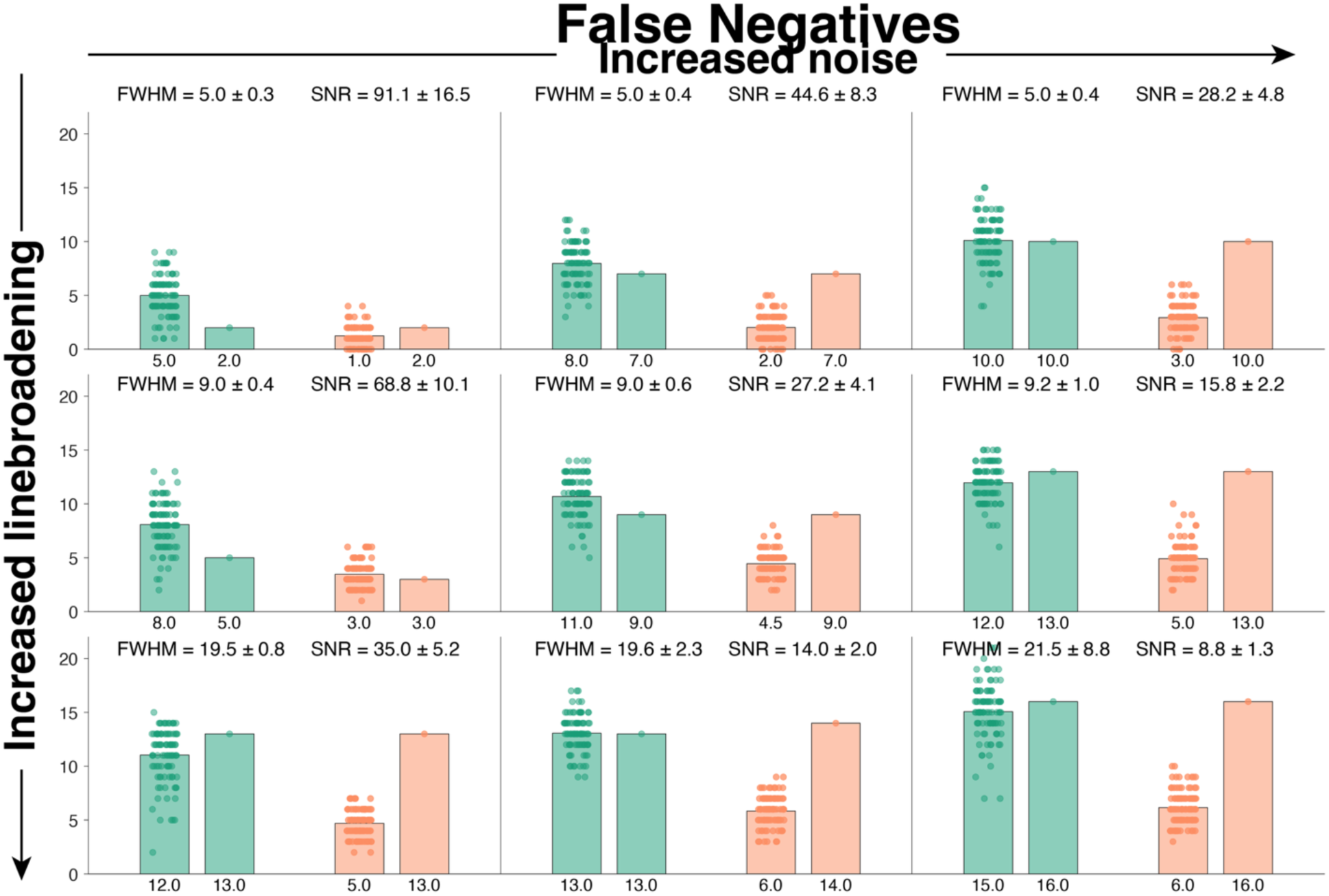
False negatives (wrongly excluded) for each of the 9 data-quality scenarios. Bars are colored by stopping condition—Max-AIC (green) and zero-Amp (orange)—with the single-spectrum optimization on the left (multiple data points) and group-wise optimization on the right (single data point). Labels at the bottom indicate the median of the distribution.

**Figure S9.**
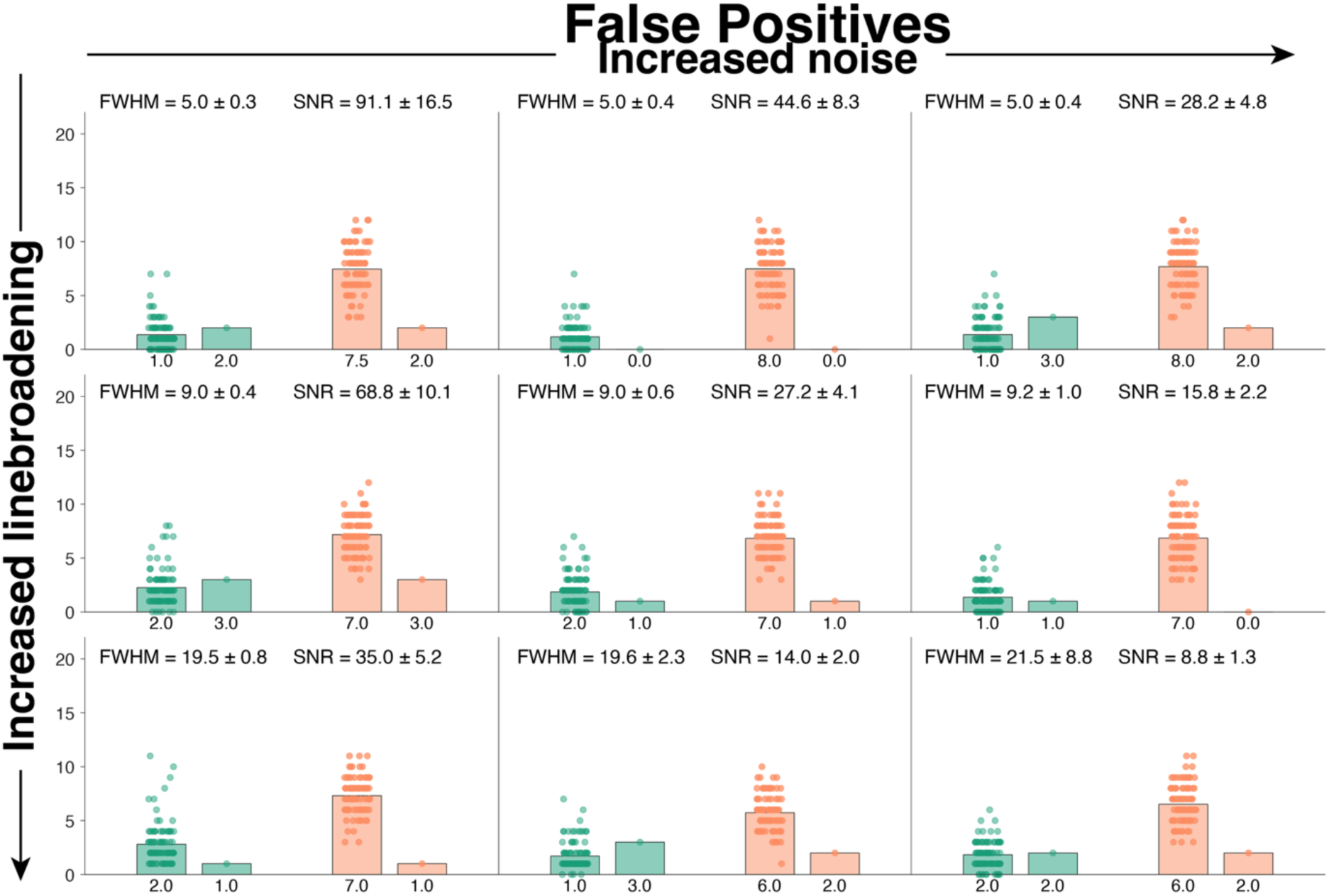
False positives (wrongly included) for each of the 9 data-quality scenarios. Bars are colored by stopping condition—Max-AIC (green) and zero-Amp (orange)—with the single-spectrum optimization on the left (multiple data points) and group-wise optimization on the right (single data point). Labels at the bottom indicate the median of the distribution.

Interestingly, lower SNR and larger FWHM adversely affected the group-wise optimizations, but the single-spectrum optimization was surprisingly more robust to the decreases in data quality.

The single-spectrum Zero-Amp optimization maintains a median 74% overlap with the ground truth basis set even for the lowest quality data. We hypothesize that there is less between-spectrum agreement in these lower-quality spectra, and this adversely affects the (median) AIC scores when they are aggregated across the entire group.

Amplitude error distributions are shown for 5 major metabolites in ***Figure S10***. While the model was comparatively robust to noise, we found that as the synthetic data’s linewidth increased, the contribution of the baseline to the model also increased, reducing the estimates of NAA and NAAG, among others. This was true even when using the correct basis set, and so represented a quirk of the precise model implementation, not specifically the algorithm itself. This was true for other metabolites, too.

**Figure S10.**
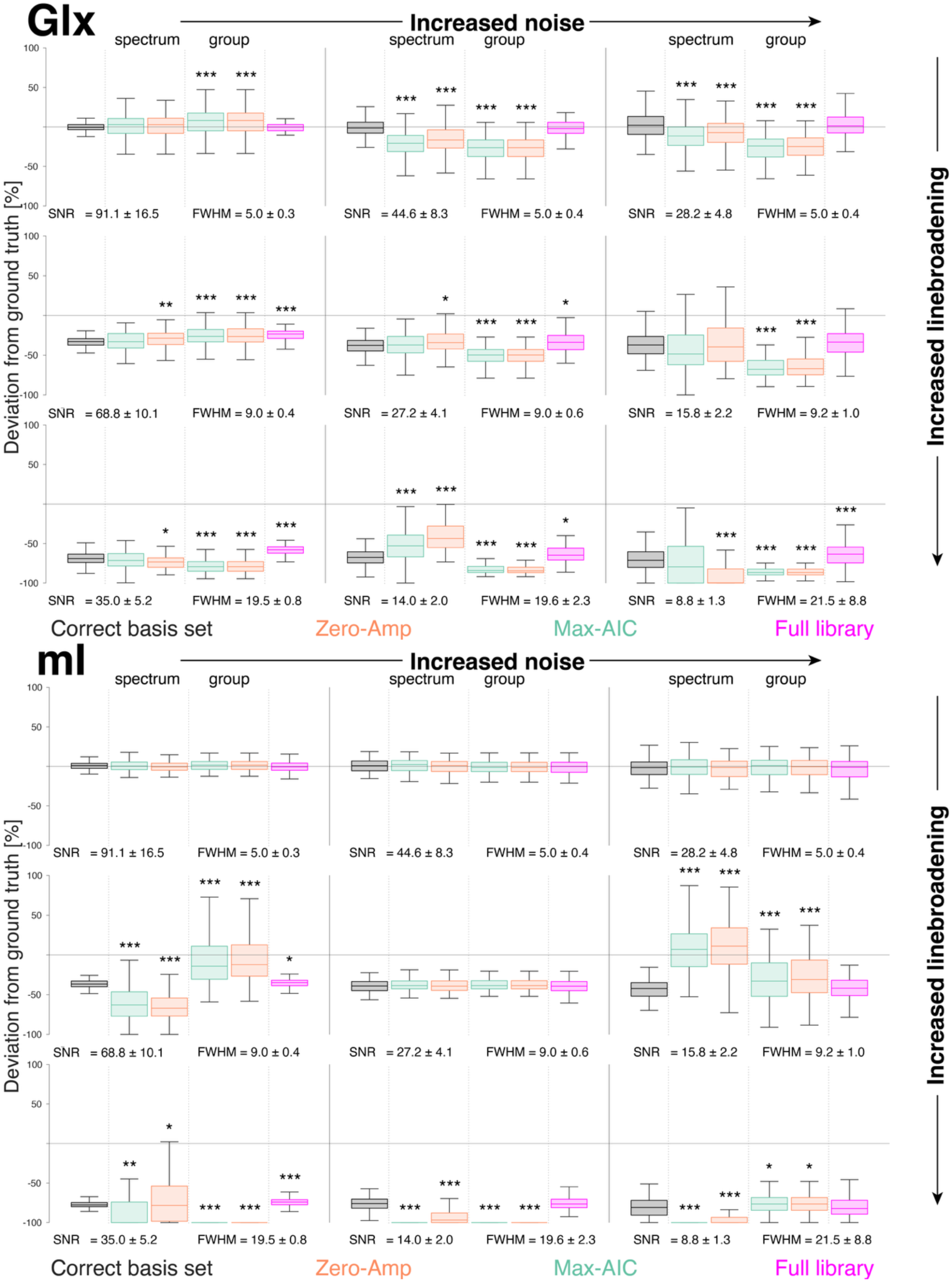
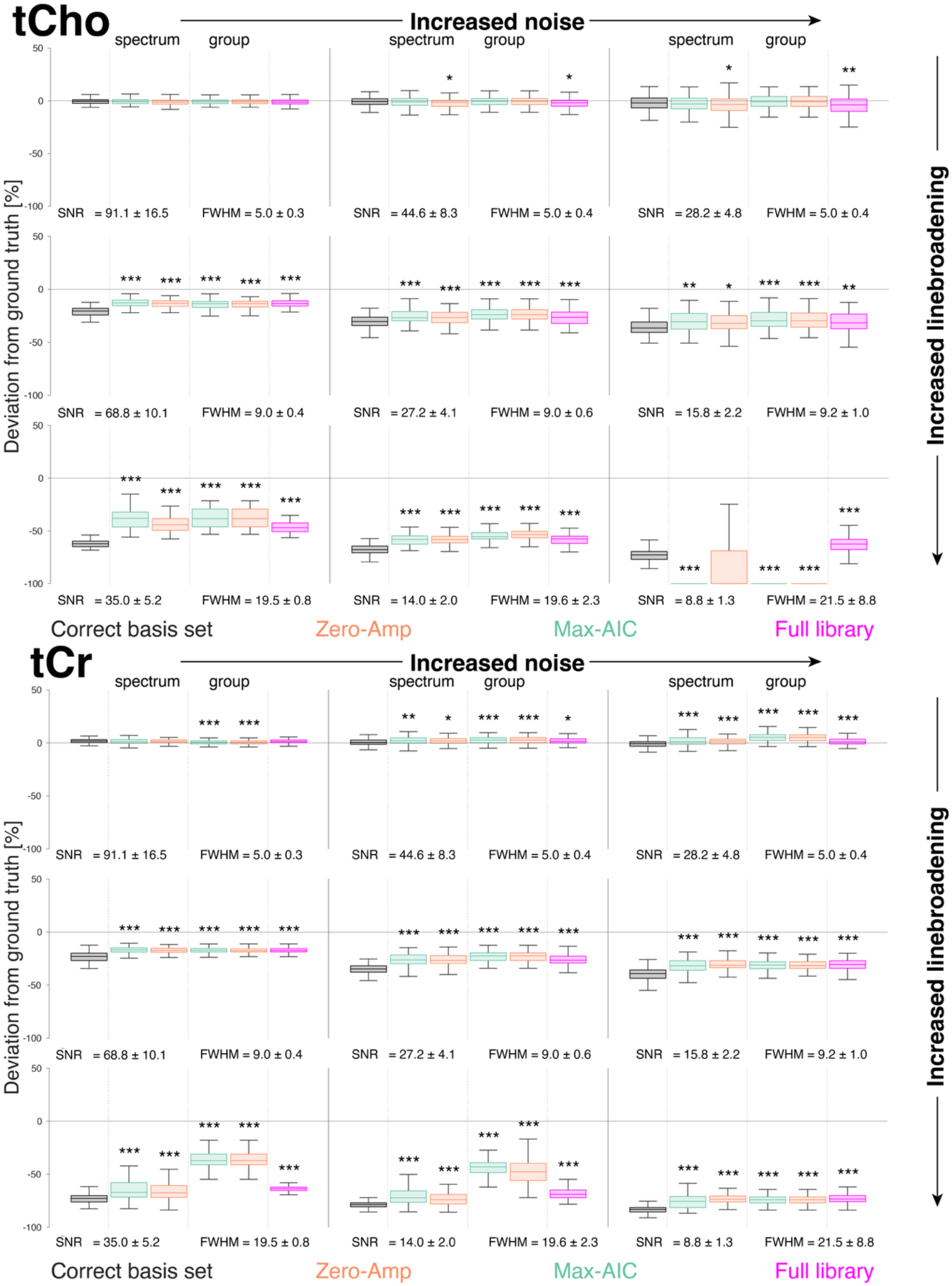
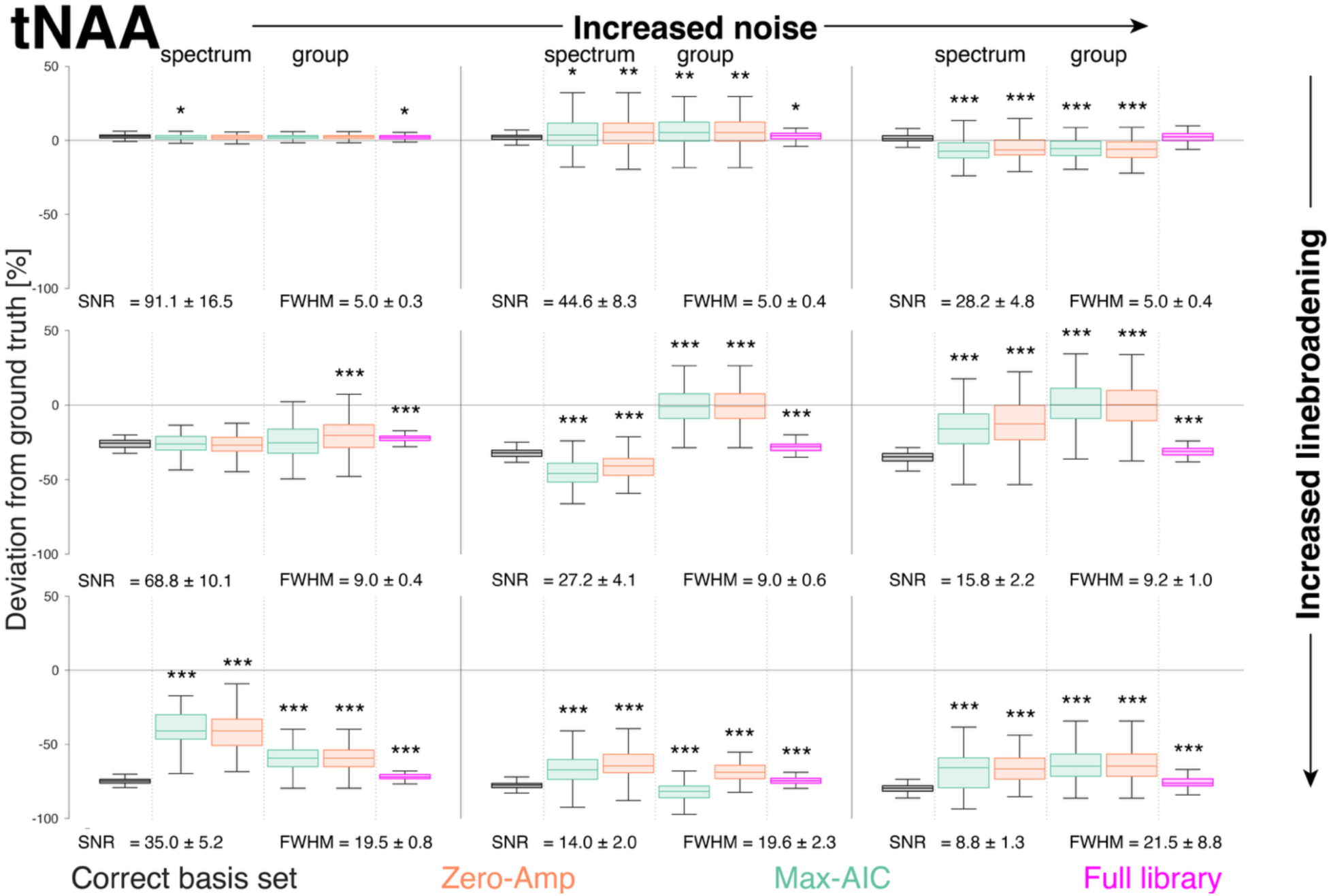
Boxplots of the percentage deviation of amplitude estimates from the simulated ground truth for tNAA, tCr, tCho, mI, and Glx. Panels represent different data quality scenarios. The results using the ground truth basis set (i.e,. including only the metabolites present in the simulation) are shown in black and the full library set in pink. The two stopping conditions— max-AIC (green) and zero-Amp (orange)—are shown in the middle sections of each panel for the single-spectrum optimization (left) and group-level optimization (right). Significant deviations from the estimates of the ground-truth basis set are marked with asterisks: “*” (0.005 ≤ p < 0.05), “**” (0.0005 ≤ p < 0.005), and “***” (p < 0.0005).

The effect of the algorithm is less clear, and results depended on the metabolite in question. Generally, the amplitude errors of the derived basis sets aligned with the ground truth for the narrowest linewidths. As linewidth increased, however, the amplitude error distributions deviated from those of the ground truth basis set and full library. Effects were metabolite-specific, but for many major metabolites (e.g., tNAA, mI, tCr, and Cho) amplitude errors from derived basis sets often had better accuracy than even the ground truth (closer to zero amplitude error) but lower precision (large distribution of errors).

## Supplementary Analysis 2: In vivo data Methods

In order to investigate how the algorithm behaved with real in vivo spectra, we utilized a large, freely available data repository^1^. Briefly, the MRS data were acquired from healthy volunteers across the age span using a 3T Philips scanner. Metabolite data were acquired using PRESS (TR/TE = 2000/30 ms; 30 x 26 x 26 mm^3^ CSO voxel; 96 transients; 2 KHz bandwidth).

Data were processed in Osprey^2^, using only the “OspreyLoad” and “OspreyProcess” modules to prepare processed spectra for input to the algorithm. A new basis set was simulated using MRSCloud^3^—appropriately representing the different sequence choice and parameters used to generate these data—but the same list of 40 metabolites was included in the library set.

Otherwise, the algorithm procedures were unchanged from the main analysis, and we ran single-spectrum and group-wise optimizations for both the maximum-AIC and zero-amplitude stopping conditions.

### Results

Figure S11 shows the AIC score trajectories for the groupwise optimization and three examples from the single-spectrum optimization. Figure 8 shows representations of how metabolites were prioritized in the single-spectrum optimization.

**Figure S11.**
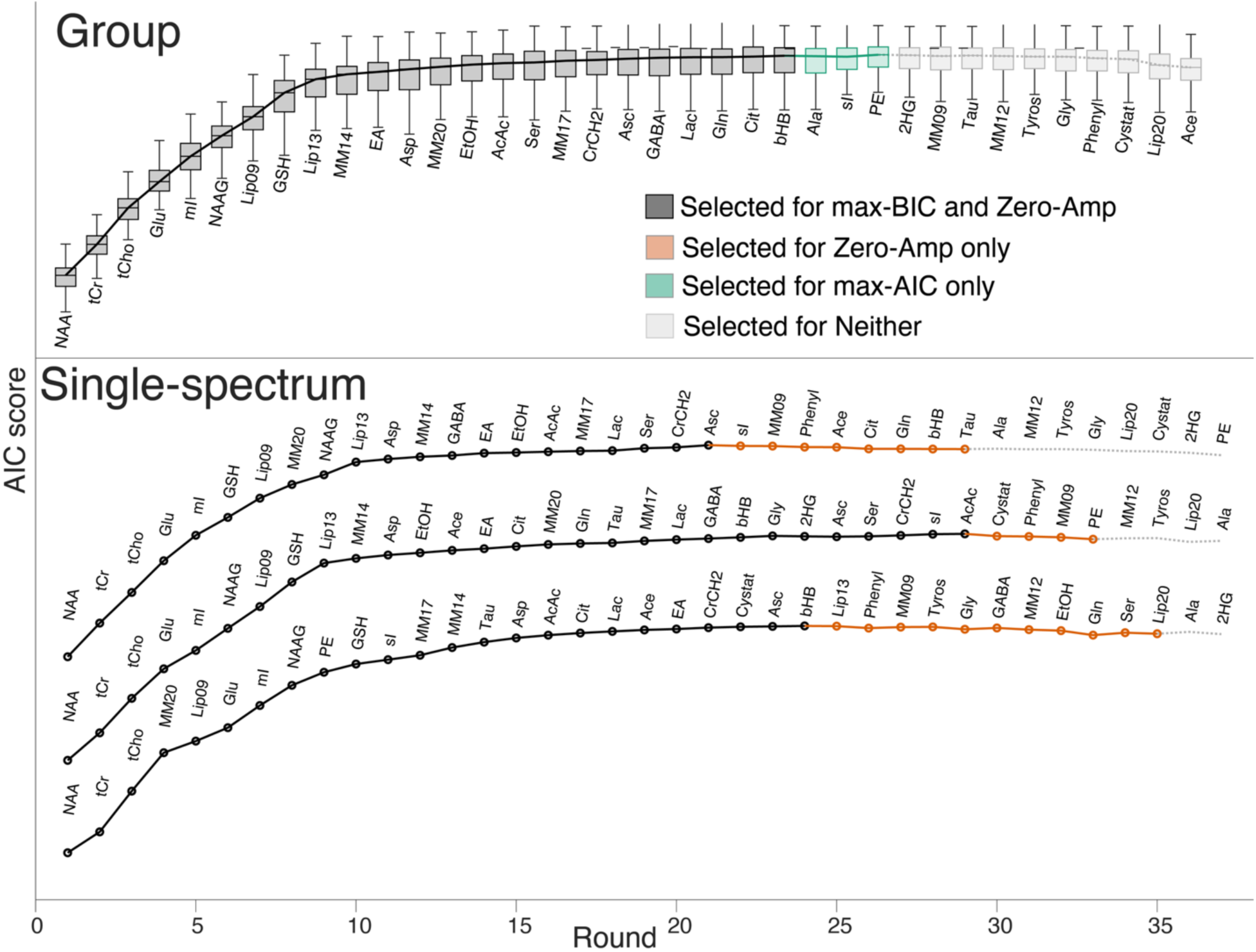
AIC trajectory derived by group-wise optimization (top) and 3 example AIC trajectories for single-spectrum optimization (bottom). Line color indicates inclusion in both stopping conditions (black, solid line), only zero-amplitude (orange, solid line), inclusion in only the max-AIC (green, solid line), and excluded from both (grey dotted line).

The most notable omissions from the group-wise-optimized basis set were MM09, MM12, and Lip20. We suspect that the model—at least, according to the AIC—can sufficiently describe these lipid and MM signals using the overlapping resonances that *were* included (Lip09, Lip13, and MM20) in synergy with specific Lorentzian linebroadening and the baseline. The benefit of these omissions is debatable and might suggest that we might need to group the lipid/MMs as we had previously done for, e.g., tCr.

In contrast to our expectations, the algorithm highly prioritized EA and excluded PE, although there may be some literature basis for this in measured concentrations^4^. EtOH and Cit were also unexpectedly included in the basis set, but this might be due to their profiles improving the modeling of aspartyl/aspartate signals at 2.5 ppm and lipid signals, respectively. This suggests that our approach is sensitive to model inaccuracies (BOTH density matrix simulations and parameterization of lipids/MMs), but without a ground truth with which to compare, these preliminary findings require further validation.

---

**Figure S12.**
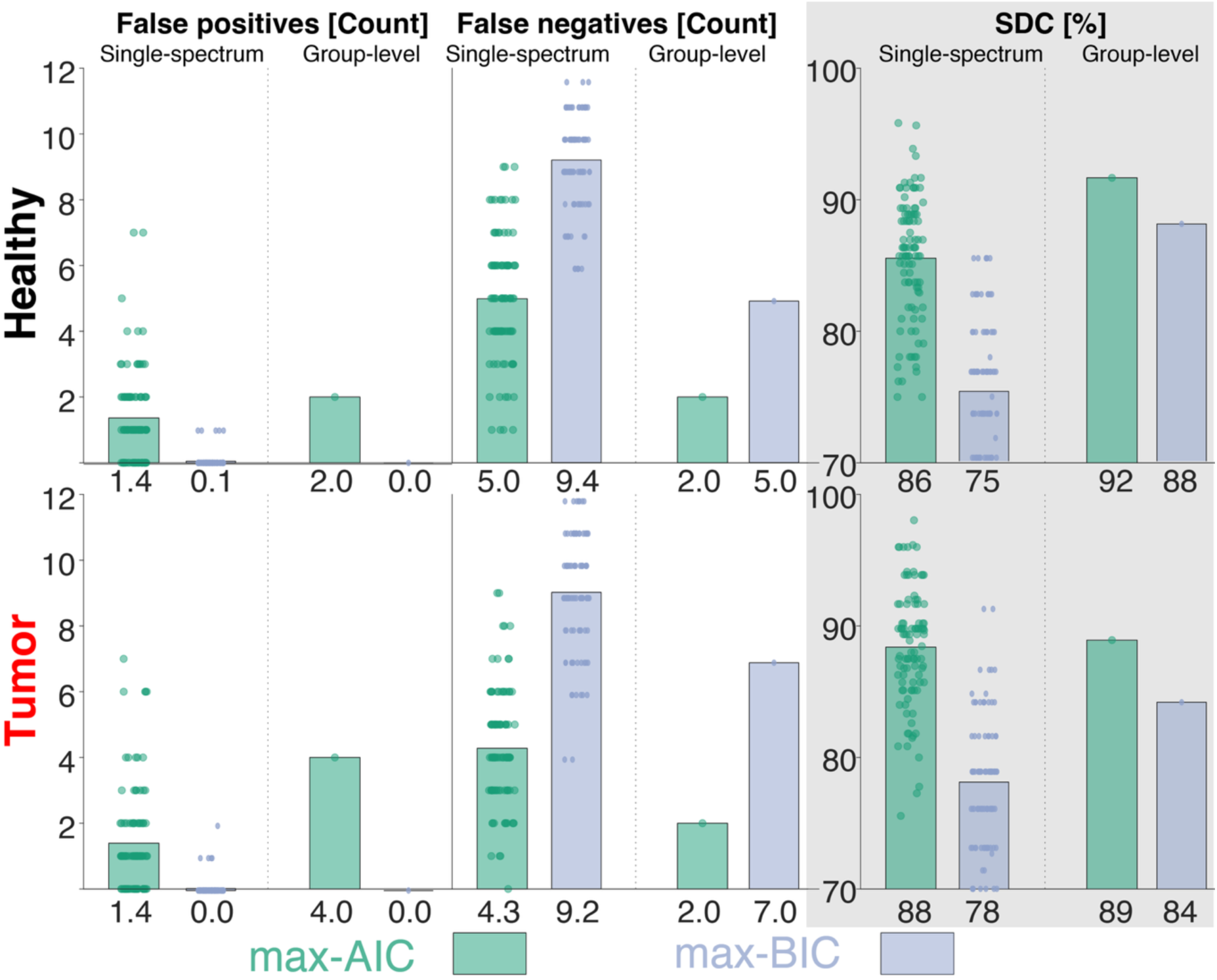
Overlap of the derived basis sets compared to the ground truth for the healthy spectra (top) and tumor spectra (bottom). Stopping conditions are color-coded: Max-AIC (green) and max-BIC (purple). From left to right, columns show the number of false positives, the number of false negatives, and the Sorensen-Dice coefficient (SDC) reported as a percentage. Within each panel, the left half shows the 100 single-spectrum results, and the right half shows the sole result of the group-wise optimization. Below each bar, the mean value is reported.

